# Segregation lift: A general mechanism for the maintenance of polygenic variation under seasonally fluctuating selection

**DOI:** 10.1101/115444

**Authors:** Meike J. Wittmann, Alan O. Bergland, Marcus W. Feldman, Paul S. Schmidt, Dmitri A. Petrov

## Abstract

Most natural populations are affected by seasonal changes in temperature, rainfall, or resource availability. Seasonally fluctuating selection could potentially make a large contribution to maintaining genetic polymorphism in populations. However, previous theory suggests that the conditions for multi-locus polymorphism are restrictive. Here we explore a more general class of models with multi-locus seasonally fluctuating selection in diploids. In these models, loci first contribute additively to a seasonal score, with a dominance parameter determining the relative contributions of heterozygous and homozygous loci. The seasonal score is then mapped to fitness via a monotonically increasing function, thereby accounting for epistasis. Using mathematical analysis and individual-based simulations, we show that stable polymorphism at many loci is possible if currently favored alleles are sufficiently dominant with respect to the additive seasonal score (but not necessarily with respect to fitness itself). This general mechanism, which we call “segregation lift”, operates for various genotype-to-fitness maps and includes the previously known mechanism of multiplicative selection with marginal overdominance as a special case. We show that segregation lift may arise naturally in situations with antagonistic pleiotropy and seasonal changes in the relative importance of traits for fitness. Segregation lift is not affected by problems of genetic load and is robust to differences in parameters across loci and seasons. Under segregation lift, loci can exhibit conspicuous seasonal allele-frequency fluctuations, but often fluctuations may also be small and hard to detect. Via segregation lift, seasonally fluctuating selection might contribute substantially to maintaining genetic variation in natural populations.

## Introduction

Ever since biologists were first able to detect population genetic variation at the molecular level, they have been puzzled by its abundance in natural populations [1]. Dispute over the underlying reasons gave rise to two scientific schools [2, 3]. Proponents of the “(neo)classical” school claim that the bulk of genetic variation is due to neutral or weakly deleterious mutations present at an equilibrium between mutation, genetic drift, and selection. The neoclassical view admits that selection may maintain alleles at intermediate frequency at some loci, but argue that such loci are exceedingly rare on a genomic scale [2]. By contrast, the “balance” school posits that substantial variation is maintained by some form of balancing selection (with some controversy over the meaning of “substantial” [3]), for example heterozygote advantage (overdominance), negative frequency-dependent selection, and spatial or temporal variability in selection pressures [4, 5].

Fifty years later, much is still open [6, 7] but the general view seems to be that (nearly) neutral mutations cause most genetic variation, with overdominance playing a relatively minor part, perhaps acting at only tens of loci per species [8–10]. A mechanism considered more common and powerful is spatial environmental heterogeneity. Temporal heterogeneity, by contrast, is believed to be of limited importance [11], despite widespread fluctuations in the strength and direction of selection, both on phenotypes [12] and on genotypes [13]. In fact, most organisms with multiple generations per year experience a particular type of temporal heterogeneity: seasonality, for example in temperature, rainfall, resource availability, but also in the abundance of predators, competitors, or parasites. Even tropical populations usually experience some seasonality [14, 15]. For example, flowering and fruiting in tropical forests is often synchronized within and even between tree species, leading to seasonal changes in food availability for animals [15]. Often, there are life-history trade-offs across seasons [16, 17]. For example, seasons with abundant resource supply might select for investment in reproduction, whereas stressful seasons may select for investment in survival. Since such life-history traits are usually polygenic, many organisms should experience seasonally fluctuating selection at a large number of loci.

With discrete generations, the fates of genotypes under temporally fluctuating selection depend on their geometric mean fitnesses over time [18]. In haploids, two alleles generally cannot coexist because one will have a higher geometric mean fitness and eventually go to fixation [18], but see [19]. In diploids, polymorphism at a single locus is stable if heterozygotes have the highest geometric mean fitness across generations (“marginal overdominance”), although in any particular generation one of the homozygotes might be fittest and increase in frequency [18, 20, 21]. However, it is not trivial to extend these results to the multi-locus case. So far, only two cases are well-understood: 1) multiplicative selection across loci, and 2) temporally fluctuating selection on a fully additive trait.

Under multiplicative selection in an infinite population with free recombination, the allele-frequency dynamics at a focal locus are independent of those at other loci. Thus, exactly as in the single-locus case, polymorphism is stable if heterozygotes have the highest geometric mean fitness. However, deviations from multiplicative selection appear to be the rule. In particular, beneficial mutations often exhibit diminishing-returns epistasis [22–24]. Additionally, there is the potential problem of genetic load. Genetic load is commonly defined as the difference between the population’s average fitness and the fitness of the fittest possible genotype. Lewontin and Hubby [1] noticed that this value can become unsustainably high if there is strong heterozygote advantage at many loci. This was a conundrum for the neoclassical school, which was worried that with high genetic load, single individuals would have to produce an astronomically large number of offspring. Others have dismissed this concern, arguing for example that selection does not generally act on all loci independently or that only relative fitness differences within the population are relevant, not fitness relative to some optimum genotype that might not even exist [25–28]. However, debate continues over whether genetic load should be an important consideration [29, 30].

The second previously studied scenario is seasonally fluctuating selection on a trait to which loci contribute additively [31, 32]. These models generally assume additivity also within loci, such that the contribution of heterozygotes is exactly intermediate between the contributions of the two homozygotes. Temporally fluctuating selection can then cause intermediate trait values to be best in the long run [33], i.e. variance in fitness is selected against and it is best to be a “jack of all trades”. This effectively is stabilizing selection on the temporal average. As such, it can generally maintain polymorphism at only one locus [34, 35], or two loci if their effect sizes are sufficiently different [36] or if they are closely linked [31]. The reason is that with multiple loci and additivity within and between loci, there are multiple genotypes with intermediate phenotypes. For two loci, for example, there is the double heterozygote (“heterozygous intermediate”) and the genotype homozygous at both loci but for alleles with opposite effects (“homozygous intermediate”). These genotypes may all have the same high fitness. However, matings between heterozygous intermediates produce a range of different genotypes, some of which are less fit than their parents. By contrast, matings between homozygous intermediates only produce new homozygous intermediates. Homozygote intermediates can therefore go to fixation and eliminate all polymorphism.

In summary, multiplicative seasonal selection is a powerful mechanism to maintain multilocus polymorphism, but the required independence across loci and the associated load call into doubt its plausibility. On the other hand, selection on additive traits can maintain polymorphism at only few loci. So far, there has been little need for further exploration because there were no clear empirical examples to challenge these theoretical results. However, part of the reason why balancing selection is so rarely observed may be that it is simply hard to detect. With recent advances in sequencing technology, a more detailed picture of genetic variation across time, space, and species is emerging, and some of the new results question the consensus that temporal heterogeneity rarely maintains variation. For instance, by sampling the same population at several time points, Bergland et al. [37] identified hundreds of loci in the genome of *Drosophila melanogaster* that exhibit strong allele frequency fluctuations in temperate populations, but are also shared with African populations of *D. melanogaster* and some even with the sister species *D. simulans,* indicating that they constitute ancient balanced polymorphisms. More generally, recent population genomic data appear to suggest that the role of balancing selection in maintaining polymorphism might be more important than previously assumed [38], and that mutation-selection-drift balance alone is not sufficient to reconcile evidence from population genomics and evidence from quantitative genetics [39]. Thus we need to reconsider the potential of temporally fluctuating selection to maintain multi-locus polymorphism.

As explained above, the conditions for multi-locus polymorphism under seasonally fluctuating selection have been examined mostly in two narrow cases. Here we examine a more general class of seasonal selection models with various forms of dominance and epistasis. Using deterministic mathematical analysis and stochastic simulations, we show that multi-locus polymorphism is possible if the currently favored allele at any time is sufficiently dominant, with dominance measured using a scale on which contributions across loci are additive. This mechanism, which we call *segregation lift,* can maintain polymorphism at a large number of loci across the genome, is robust to many model perturbations, and does not require single individuals to have too many offspring. Depending on the parameter values, allele-frequency fluctuations can be large and readily detectable, or subtle and hard to discern.

## Basic model

We consider a diploid, randomly mating population in a seasonally fluctuating environment. Specifically, we assume a yearly cycle with g generations of winter followed by *g* generations of summer (robustness to asymmetry is explored below). The genome consists of *L* unlinked loci with two alleles each: one summer-favored and one winter-favored allele. For a given multi-locus genotype, let *n_s_* and *n_w_* be the number of loci homozygous for the summer and winter allele, respectively, and *n_het_* the number of heterozygous loci, with *n_s_* + *n_w_* + *n_het_* = *L*.

In the basic model, loci are interchangeable in their effects (see Stochastic simulations for a more general model) and the fitness of a multi-locus genotype can be computed as a function of *n_s_*, *n_w_* and *n_het_*. In the simplest case, fitness depends only on *n_s_* + 0.5 *n_het_* in summer and *n_w_* + 0.5 *n_het_* in winter, i.e. half the number of currently favored allele copies. To allow for dominance effects, we generalize this simple scenario and assume that fitness in summer depends on the *summer score*

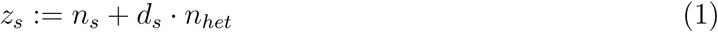

and fitness in winter depends on the *winter score*

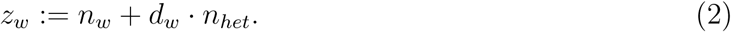

The parameters *d_s_* and *d_w_* quantify the dominance of the currently favored allele in summer and winter, not with respect to fitness, but with respect to the seasonal scores *z_s_* and *z_w_*. Because we are interested in whether temporally fluctuating selection can maintain polymorphism in the absence of other stabilizing mechanisms, we only consider values of *d_s_* and *d_w_* between 0 and 1, and do not allow values larger than one, which would correspond to standard heterozygote advantage.

The relationship between the seasonal score *z* (*z* = *z_s_* in summer and *z* = *z_w_* in winter), and fitness, *w,* is given by a monotonically increasing fitness function *w*(*z*). This function specifies the strength of selection and accounts for epistasis. With discrete generations, the allele-frequency dynamics at a focal locus are driven by the relative fitnesses of the three possible genotypes at that locus, for example the ratio of the fitness of homozygotes and heterozygotes. We say that there is no epistasis if these ratios and thus the strength of selection are independent of the number of other loci and their contributions to *z*. This is the case when fitness is multiplicative across loci:

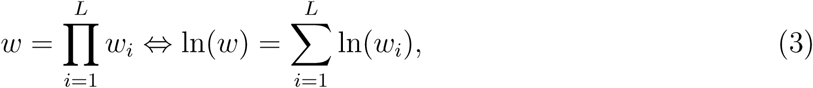

where *w_i_* is the fitness value at locus *i.* In our model, this is achieved by setting *w*(*z*) = exp(*z*) because then Eq. (3) is fulfilled with *w_i_* = exp(1) if locus *i* is homozygous for the currently favored allele, *w_i_* = exp(*d_s_*) or *w_i_* = exp(*d_w_*) if it is heterozygous, and *w_i_* = exp(0) = 1 if it is homozygous for the currently disfavored allele. With *υ*(*z*): = ln(*w*(*z*)), the multiplicative model is characterized by υ″(*z*) = 0. We therefore use the second derivative of the logarithm of fitness *υ″*(*z*) as a measure of epistasis (see [40] for a similar definition of epistasis). Under positive or synergistic epistasis (*υ″*(*z*) *>* 0), the logarithm of fitness increases faster than linearly with *z* and thus selection at a focal locus increases in strength with increasing contribution of the other loci to *z*. By contrast under negative or diminishing-returns epistasis (υ″(*z*) < 0), selection at a focal locus becomes weaker with increasing contribution of other loci to *z*. We focus on two classes of fitness functions (Fig. 1). The first is of the form

**Figure 1:**
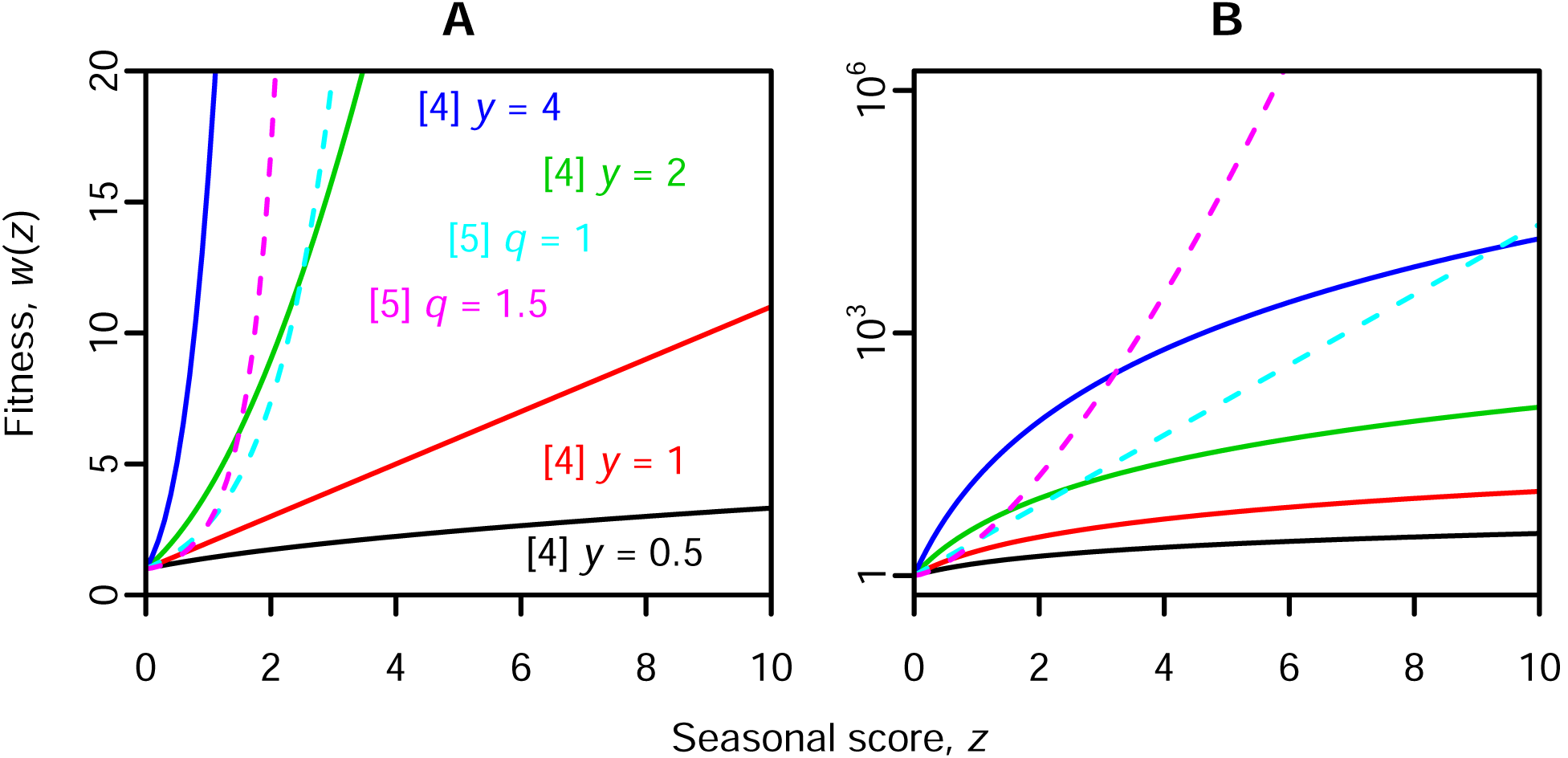
Examples for fitness functions generated by Eq. (4) or Eq. (5) with various parameters. In (A), fitness is shown on a linear scale, in (B) on a logarithmic scale.

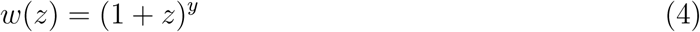

with a positive parameter *y* for the strength of selection. Although for *y >* 1 in Eq. (4), fitness increases faster than linearly with increasing *z* (Fig. 1 A), transformation to the logarithmic scale reveals that epistasis is negative for all *y* (Fig. 1 B). The second class of fitness functions is of the form

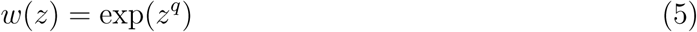

with *q* ≥ 1. This function reduces to the multiplicative model with *q* =1 (cyan lines in Fig. 1) and has positive epistasis with q > 1 (e.g. magenta lines in Fig. 1).

In summary, fitness is computed in two steps. The first maps the multi-locus genotype onto a seasonal score *z* to which loci contribute additively, essentially a generalized counter of the number of favored alleles (Eq. (1), Eq. (2)), and the second maps *z* to fitness (Fig. 1). This two-step process disentangles dominance (first step) and epistasis (second step).

In our model, there is antagonistic pleiotropy in the sense that genotypes with many summer alleles have a high summer score but a low winter score and vice versa. Previous theoretical studies suggest that antagonistic pleiotropy is most likely to maintain polymorphism if for each trait affected by a locus the respective beneficial allele is dominant [41, 42]. Such “reversal of dominance” also facilitates the maintenance of polymorphism in single-locus models for temporally fluctuating selection [43]. Hypothesizing that reversal of dominance would also help to maintain polymorphism under multi-locus temporally fluctuating selection, we assume that *d_s_* and *d_w_* in Eq. (1) and Eq. (2) take the same value, *d,* which means that dominance switches between seasons (see Stochastic simulations for a more general model). For d < 0.5, we have “deleterious reversal of dominance” [42] and the currently favored allele is always recessive, whereas for d > 0.5, we have “beneficial reversal of dominance” [42] and the currently favored allele is always dominant. If d = 0.5, the seasonal score z is additive not just between loci, but also within loci. Importantly, the value of d also determines the relative fitness of “heterozygous intermediates”, multi-locus genotypes with the same number of summer and winter alleles and at least some heterozygous loci, compared to “homozygous intermediates”, which also have the same number of summer and winter alleles but are fully homozygous (Fig. 2). For *d* < 0.5, heterozygous intermediates have a lower seasonal score, *z*, and therefore a lower fitness in both seasons than homozygous intermediates. For *d* = 0.5, heterozygous and homozygous intermediates have the same score and fitness. Finally, for *d* > 0.5 heterozygous intermediates have a higher score and fitness.

**Figure 2:**
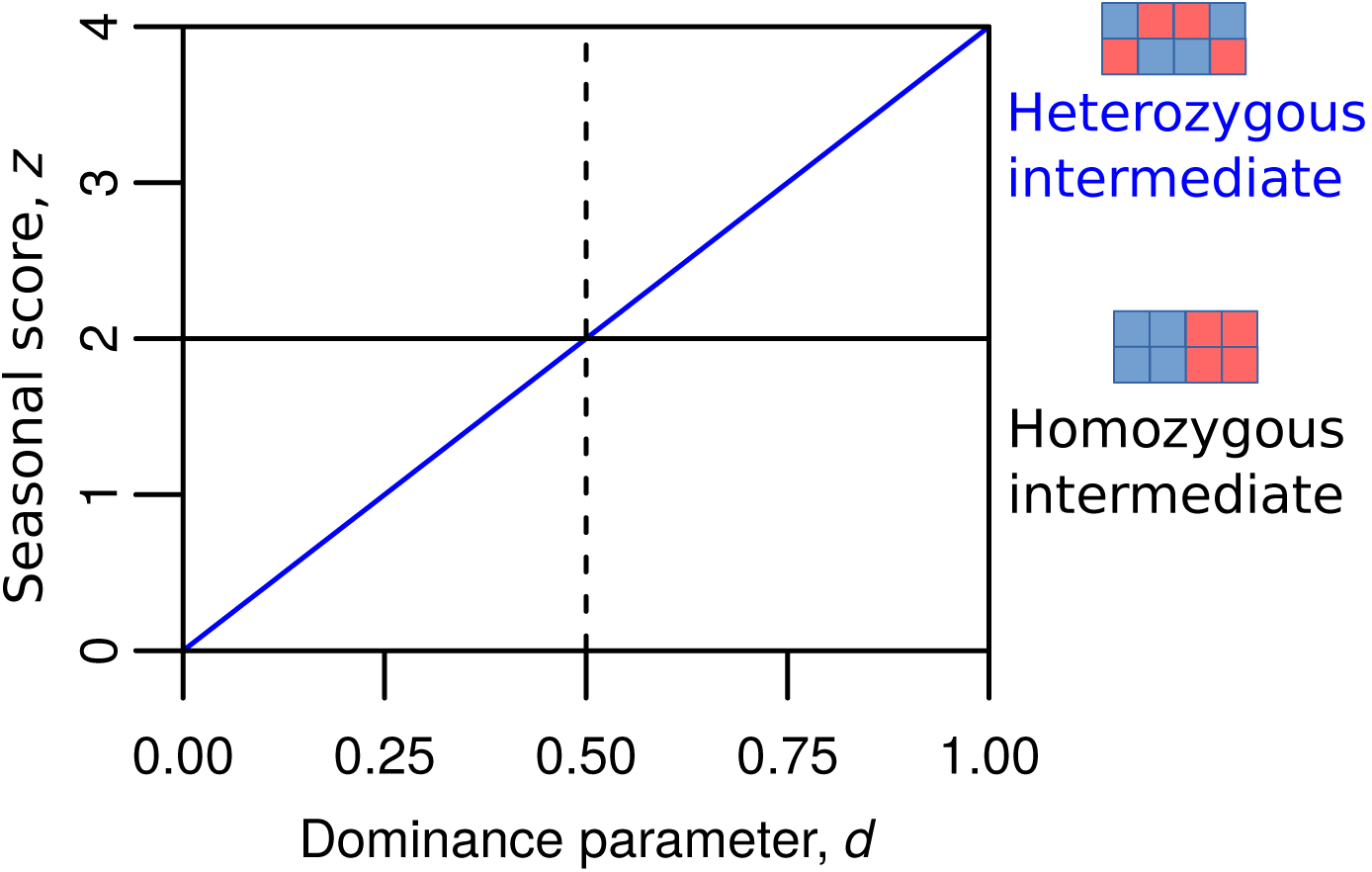
Values of the seasonal score, *z*, as a function of the dominance parameter, *d,* for two example four-locus genotypes: a heterozygous intermediate (blue line) and a homozygous intermediate (black solid line).

Reversal of dominance between seasons can arise naturally in situations with antagonistic pleiotropy for two traits. In the example in Fig. 3, *d* > 0.5 in both seasons although the effects of the different alleles on the two traits remain constant throughout the year. Seasonal changes are required only in the relative importance of traits, which appears common in natural populations. For example, in many species, starvation tolerance might be more important in winter and fecundity more important under abundant resource supply in summer. Note that this particular example also requires changes (though not necessarily a reversal, see Fig. S1 A) in dominance with respect to the pleiotropic effects of a locus on the two traits. On the metabolic level, such changes in dominance may arise in branched enzyme pathways with saturation or feedbacks [44], but it is unclear how common such situations are. Alternatively, seasonal reversal of dominance could be mediated by seasonal reversal in gene expression. In *Drosophila melanogaster,* for instance, many genes exhibit reversal of dominance in gene expression between different environmental conditions [45].

**Figure 3:**
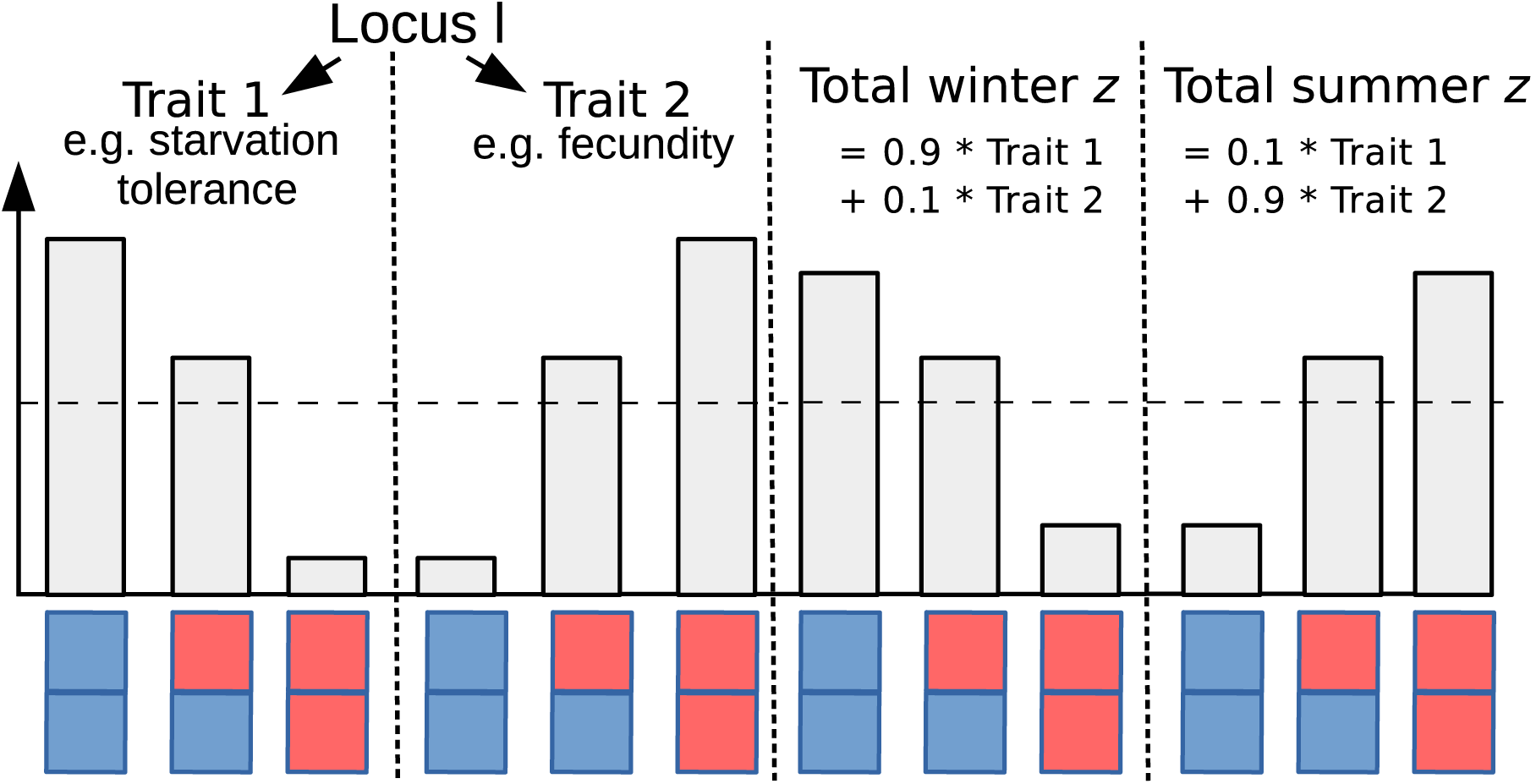
Potential mechanistic underpinning for beneficial reversal of dominance. There is antagonistic pleiotropy for two traits and the seasonal scores for winter and summer are computed as weighted averages of trait 1 and trait 2, with the relative importance of the two traits switching between seasons. The dashed line indicates the average of the two homozygote traits. If the heterozygotes are closer to the fitter homozygote with respect to both trait 1 and trait 2, there is a beneficial reversal of dominance at the level of the seasonal score, z. See Fig. S1 for alternative scenarios.

Interestingly, because *d* measures dominance not at the scale of fitness but at the scale of the seasonal score, *z,* reversal of dominance for fitness is neither sufficient nor necessary for *d* > 0.5. To see this, consider the counterexamples in Fig. 4 where we compare the genotypes at a focal locus in a common genetic background. If the fitness function is concave, the fitness of a heterozygote can be closer to the fitter homozygote even for *d* < 0.5 (Fig. 4 A). On the other hand, if the fitness function is convex, the fitness of heterozygotes can be closer to the less fit homozygote even for *d* > 0.5 (Fig. 4 B).

**Figure 4:**
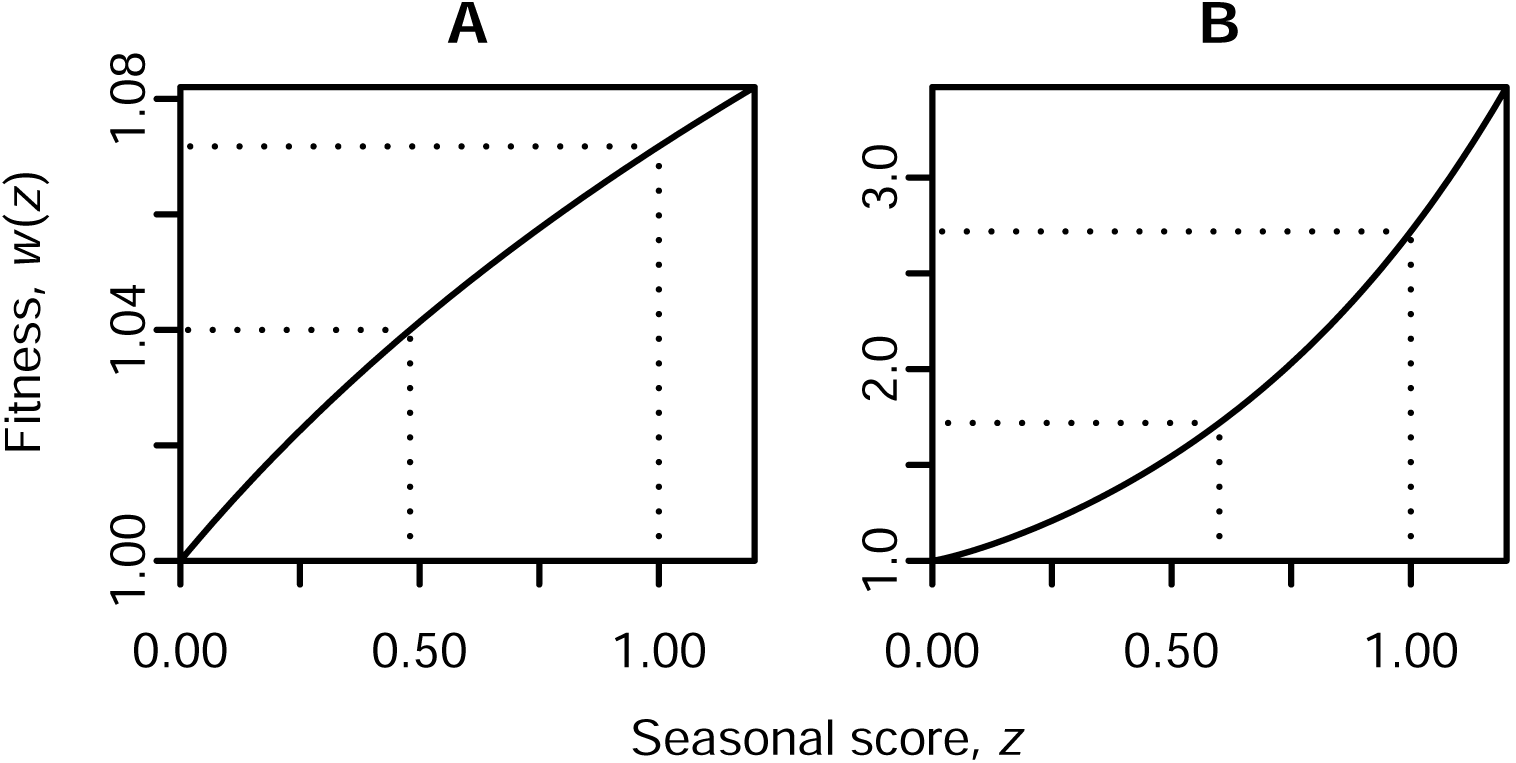
Examples illustrating why a heterozygote fitness closer to the fitter homozygote is neither a sufficient (A) nor a necessary condition (B) for *d >* 0.5. It is assumed that the genetic background is the same for all genotypes and, for simplicity, that it makes contribution 0 to *z*. The dotted lines illustrate the mapping between *z* = *d* and *z* = 1 and the respective fitnesses. A) Eq. (4) with *y* = 0.1. B) Eq. (5) with *q* = 1.2.

## Deterministic analysis

In this section, we assume that population size is so large that genetic drift does not play a role. We also assume that mutations are rare enough that the allele-frequency dynamics will equilibrate before a new mutation arises at one of the *L* loci. This simple deterministic framework allows us to develop an intuitive understanding of the conditions for stable polymorphisms for various genotype-to-fitness maps. The intuitions developed here will then be checked and extended with stochastic simulations in the next section.

We will first confirm that the conditions under which seasonally fluctuating selection can maintain polymorphism are restrictive when contributions to the seasonal score z are additive within loci (*d* = *d_s_* = *d_w_* = 0.5 in Eq. (1) and Eq. (2)). Then *z_s_* + *z_w_* = *L* for all possible genotypes and the mean *z* over time is *z*\* = *L*/2. The long term success of a genotype depends on its geometric mean fitness, or equivalently on the arithmetic mean of the logarithm of fitness, *υ*(*z*). Jensen’s inequality or simple geometric considerations (Fig. 5) tell us that the arithmetic mean of *υ*(*z_s_*) and *υ*(*z_w_*) for a given genotype will be smaller than or equal to *υ*(*z*\*) if *υ″ <* 0 everywhere (Fig. 5 A), equal to *υ*(*z*\*) if *υ*″ = 0 (Fig. 5 B), and larger than or equal to *υ*(*z*\*) if *υ" >* 0 (Fig. 5 C).

**Figure 5:**
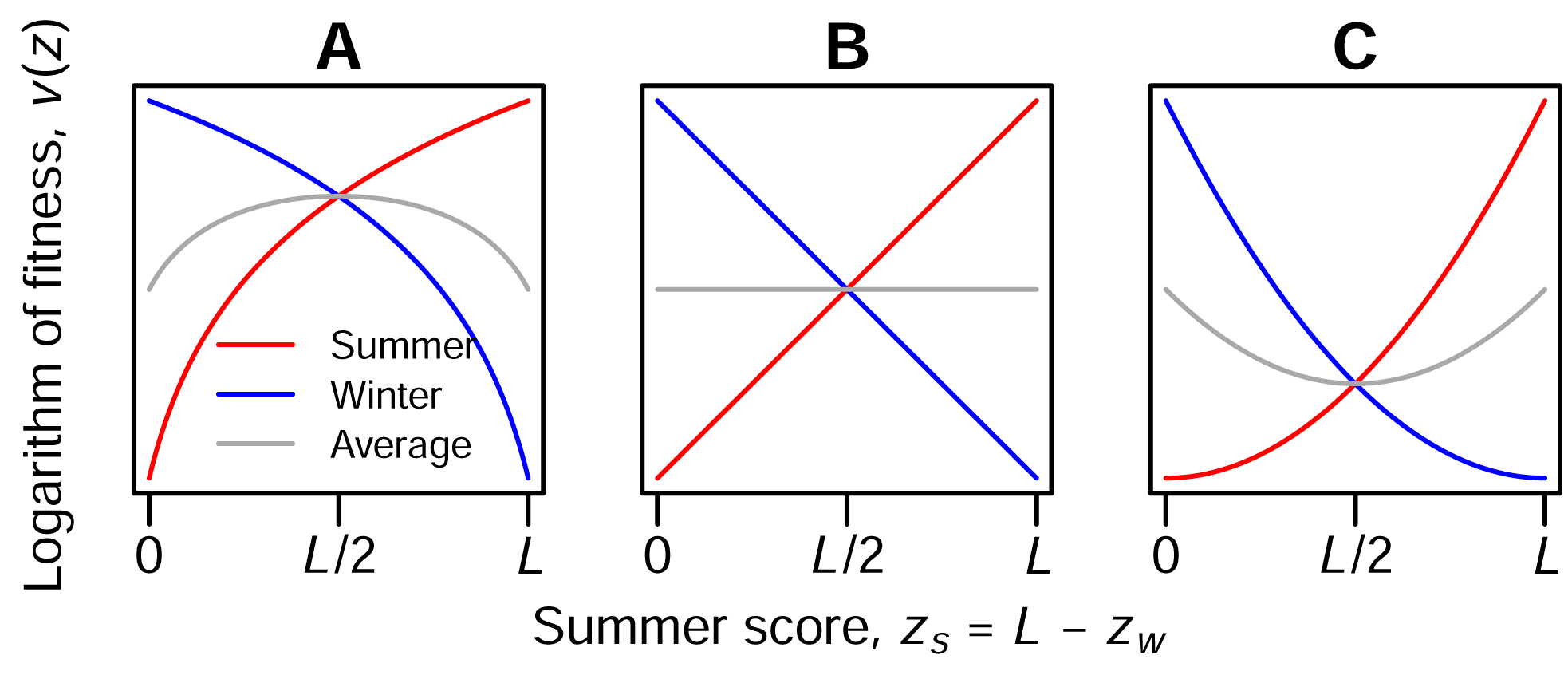
The logarithm of summer fitness (red) and winter fitness (blue) and the average logarithm of fitness (gray) as a function of a genotype’s summer score, *z_s_*. Assuming *d* = 0.5, the winter score is *z_w_* = *L – z_s_*, leading to the mirror symmetry around L/2. If the fitness function is concave on the logarithmic scale, intermediate types have the highest average log-fitness (A), if the fitness function is log-linear, then all types have the same average log-fitness (B), and if the fitness function is convex on a logarithmic scale, extreme types have the highest average log-fitness and thus the highest geometric mean fitness (C).

The inter-annual allele-frequency dynamics (e.g. from summer to summer or from winter to winter) with multiplicative fitness (*υ*″ = 0) and *d* = 0.5 are thus neutral. No balancing selection emerges. With positive epistasis (*υ*″ > 0), extreme types with either only summer alleles or only winter alleles have the highest geometric mean fitness. Therefore, the population ends up in a state where all loci are fixed for the summer allele or all for the winter allele. With negative epistasis (*υ″ <* 0), the genotypes with the highest geometric mean fitness are those with the same number of summer and winter alleles and thus *z_s_* = *z_w_*. There are always some genotypes heterozygous at one or more loci that fulfill this condition (heterozygous intermediates, see Fig. 2). For an even number of loci, *z_s_* = *z_w_* is also true for genotypes homozygous for the summer allele at half of the loci and homozygous for the winter allele at the other half (homozygous intermediates). When one of the homozygous intermediates fixes in the population it cannot be invaded by any mutant starting at small frequency (under the assumptions of the deterministic model, see Appendix S1 for a detailed proof), and all polymorphism is eliminated. For an odd number of loci, homozygous intermediates do not exist and some polymorphism may be maintained, at least at one locus. This case, which we will examine in more detail below, appears to be the only way in which seasonally fluctuating selection can maintain polymorphism under additivity (*d* = 0.5).

Next we explore whether deviations from additivity (*d* ≠ 0.5) can facilitate multi-locus polymorphism. Under multiplicative selection, i.e. without epistasis, the conditions for polymorphism at one locus are not affected by the dynamics at other loci. Thus given the fitness values for individual loci (exp(*d*) for heterozygotes, exp(1) and exp(0) for currently favored and disfavored homozygotes, respectively, see text below Eq. (3)), we can conclude that polymorphism is possible if

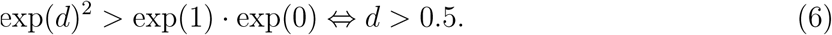

That is, there must be a beneficial reversal of dominance with respect to z such that at any time the currently favored allele is dominant.

Now we explore whether such a beneficial reversal of dominance can also maintain polymorphism in the presence of epistasis. In each case, a necessary condition for polymorphism is that a population fixed for the fittest possible fully homozygous genotype can be invaded by mutants. As we have seen above, with synergistic epistasis (*υ″ >* 0) there are two fully homozygous genotypes with maximum fitness, the one with the summer allele at all loci and the one with the winter allele at all loci. In both cases, the resident type has score *L* in one season and score 0 in the other season, whereas mutants differing in one position have scores *L* – 1 + *d* and *d.* For mutants to invade, we thus need

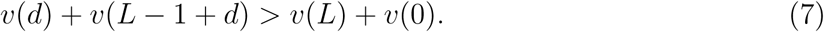

Using our example class of fitness functions with positive epistasis, Eq. (5) with *q >* 1, we thus obtain the condition

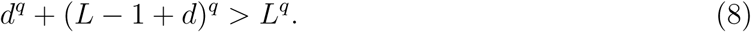

The critical value of *d*, *d_crit_*, at which extreme types become invasible, satisfies

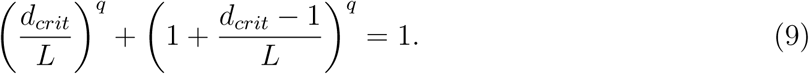

For *q* = 2, *d_crit_* takes values 0.707, 0.954, and 0.995, with 1 locus, 10 and 100 loci, respectively. For *q* > 1 in general, *d_crit_* approaches 1 as the number of loci increases. To see this, note first that the condition in Eq. (9) is always fulfilled for *d* =1 and thus *d_crit_* ≤ 1. Thus, as the number of loci, *L*, goes to infinity (*d_crit_* – 1)/*L* becomes small and we can approximate the second term on the left hand side of Eq. (9) by a Taylor expansion around 1 to obtain

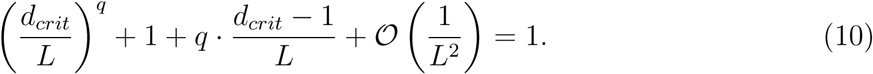

Multiplying both sides by *L* and letting *L* go to infinity, we can conclude that *d_crit_* is approximately 1 if the number of loci is large. Thus for fitness functions of the type in Eq. (5) with positive epistasis, seasonally fluctuating selection can in principle maintain polymorphism at many loci, but the respective favored allele would have to be almost completely dominant, requiring large seasonal changes in dominance.

With diminishing-returns epistasis (*υ*″ < 0), the fittest possible fully homozygous genotype carries the summer allele at half of the loci and the winter allele at the other half of the loci (assuming an even number of loci). Thus a necessary condition for polymorphism is that this homozygous intermediate type can be invaded by mutants. The resident type has score *L*/2 in both seasons whereas mutants differing in one position have scores *L*/2 + *d* and *L*/2 — 1 + *d*.

Thus the resulting necessary condition for polymorphism is

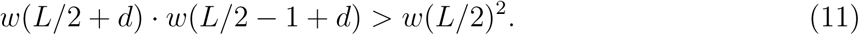

Again, this condition is always fulfilled for *d* = 1. For fitness functions of the form Eq. (4) with any exponent *y*, the critical dominance coefficient *d_crit_* at which homozygous intermediates become invasible satisfies

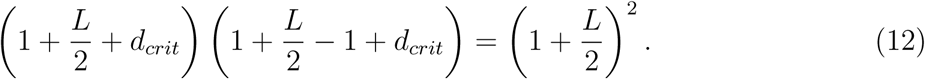

This quadratic equation has a negative solution, which is not relevant for our model, and a positive solution

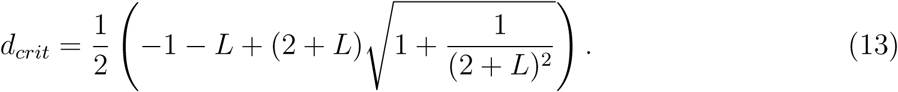

From Eq. (13), *d_crit_* decreases as *L* increases and approaches 0.5 as *L* goes to infinity. The intuition here is that the second derivative of the logarithm of fitness *υ"*(*z*) = –*y*(1 + *z*)^−2^ decreases with increasing *z*. Therefore, for large *L*, epistasis around the intermediate type with *z* = *L*/2 is weak and the conditions for polymorphism approach those without epistasis. In other words, with increasing *L*, selection against temporal variation around the intermediate type becomes weaker and a smaller change in dominance is sufficient to overcome it.

Our results so far suggest that for a broad class of fitness functions seasonally fluctuating selection can maintain polymorphism if in both seasons the respective favored allele is sufficiently dominant. We call this mechanism *segregation lift* because it is based on a positive aspect of two alleles segregating at the same locus, as opposed to the negative aspect of segregation load. However, the preceding analysis does not tell us whether polymorphism will be maintained at all loci, or just one or a few of them. Also, it is still unclear how efficient segregation lift is at maintaining multi-locus polymorphism in finite populations with genetic drift and recurrent mutations and whether genetic load is a problem. To address these questions, we now turn to stochastic simulations.

## Stochastic simulations

We use Wright-Fisher type individual-based forward simulations (see Appendix S2.1 for details). That is, for every individual in a generation independently, two individuals are sampled as parents in proportion to their fitnesses. We focus on diminishing-returns fitness functions of type Eq. (4) both because diminishing-returns epistasis is common [e.g. 22–24] and because the above theoretical arguments suggest that it is more conducive to multi-locus polymorphism than synergistic epistasis. Specifically, the critical dominance parameter, *d_crit_,* for diminishing-returns epistasis in Eq. (13) is generally smaller than the one for synergistic epistasis in Eq. (9).

Additional parameters in the stochastic simulations are the symmetric mutation probability *μ* per allele copy per generation and the population size *N*. We generally keep population size constant over the year, but we also run supplementary simulations with seasonal changes in population size. Table 1 gives an overview of the model parameters and the ranges explored. In most simulated scenarios, selection and dominance effects are strong relative to mutation (*w'*(*z*) and *d* – 0.5 are much larger than the mutation rate *μ*). Natural populations are certainly often larger and mutation rates smaller than the values used here. However, since many population genetic processes depend only on the product *Νμ* [e.g. 46], large populations with small mutation rates may be well approximated by computationally more manageable smaller populations with larger mutation rates.

**Table 1:** Overview of model parameters and the ranges explored in the simulations.

| Parameter | Explanation | Range explored |
| --- | --- | --- |
| $g$ | Number of generations per season; a year has $2g$ generations | 1–20 |
| $N$ | Population size | 100–10,000 |
| $L$ | Number of loci | 1–500 |
| $\mu$ | Mutation probability per allele copy per generation | $10^{-6} - 10^{-4}$ |
| $d$ | Dominance parameter | 0–1 |
| $y$ | Exponent of the fitness function Eq. (4) | 0.5–4 |

In addition to the basic model, we designed a “capped” model to assess the relevance of genetic load. In this model, individuals can have at most 10 offspring. While assembling the offspring generation from the parent generation, we track how many offspring a parent individual has already produced. Once this number reaches 10, the fitness of this individual is set to 0 so that it cannot be drawn as a parent again.

From the simulation output, we estimate an “effective strength of balancing selection” (see Methods and Appendix S2.2), which tells us whether and how fast a rare allele increases in frequency over a full yearly cycle. As expected from the above theoretical arguments, additive contributions within loci (*d* = 0.5) are not conducive to multi-locus polymorphism (Fig. 6). For even numbers of loci, i.e. whenever haplotypes having an equal number of summer and winter alleles are possible (homozygous intermediates), the effective strength of balancing selection estimated from the simulations is negative, indicating that rare alleles tend to become even rarer. For small odd numbers of loci, the effective strength of balancing selection is positive, but only one or two loci at a time fluctuate at intermediate frequency (Fig. S3). As the number of loci increases, the effective strength of balancing selection eventually becomes negative even for odd numbers, decreases overall in absolute value, and finally approaches zero (effective neutrality) from below (Fig. 6). This behavior is independent of the exponent, *y,* of the fitness function Eq. (4). Also as expected, effective balancing selection (see Methods) emerges if the dominance parameter, *d,* is larger than a certain critical value, which decreases with the number of loci and is only weakly influenced by mutation rate (Fig. 7).

**Figure 6:**
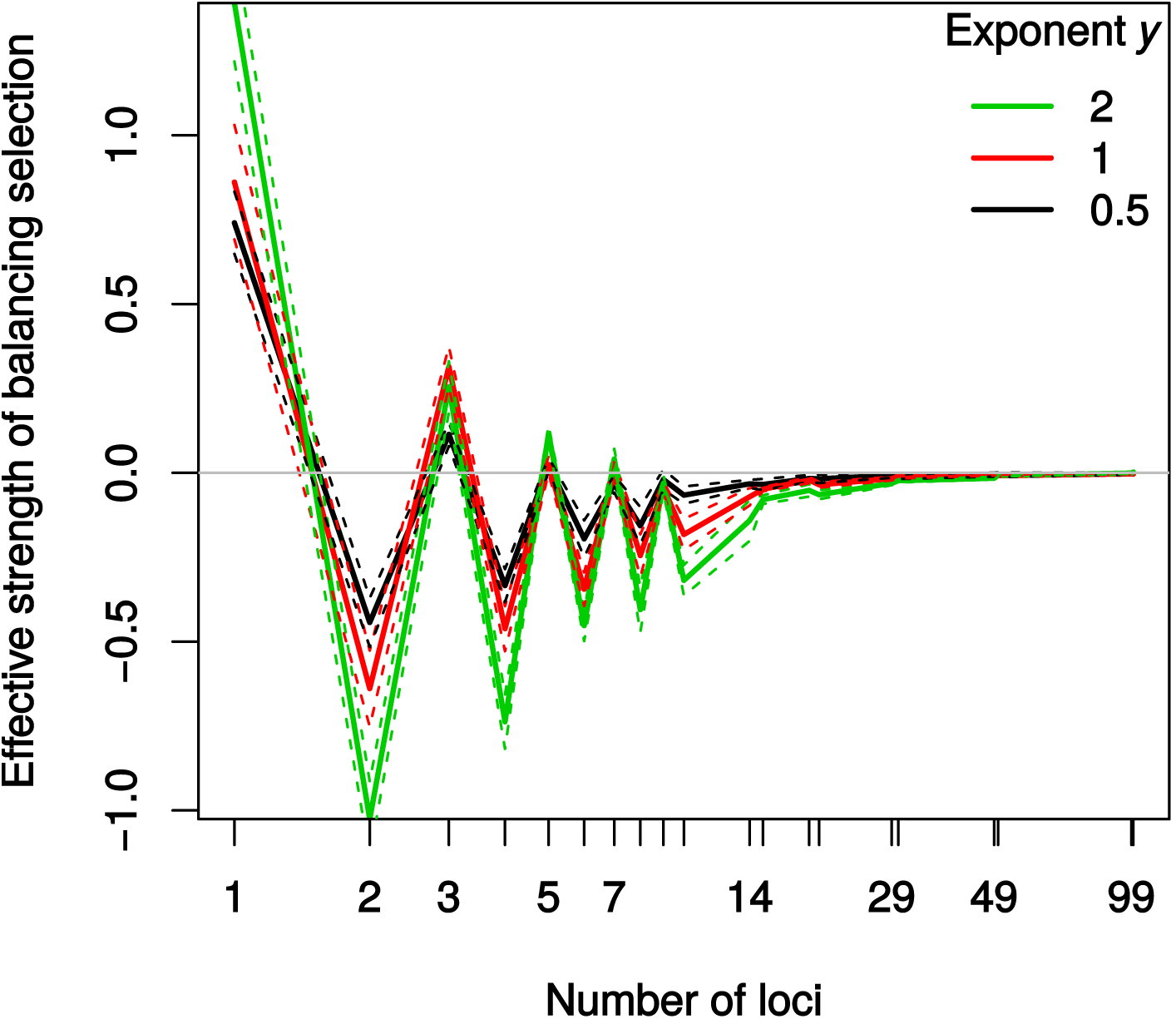
Effective strength of balancing selection (*b_e_* in Eq. (14) in Methods), in the additive case (*d* = 0.5) as a function of the number of loci. Solid lines indicate means and dashed lines indicate means ± two standard errors. Simulations were always run for successive odd and even numbers. *N* = 1000,*g* = 15, *μ* = 0.0001.

**Figure 7:**
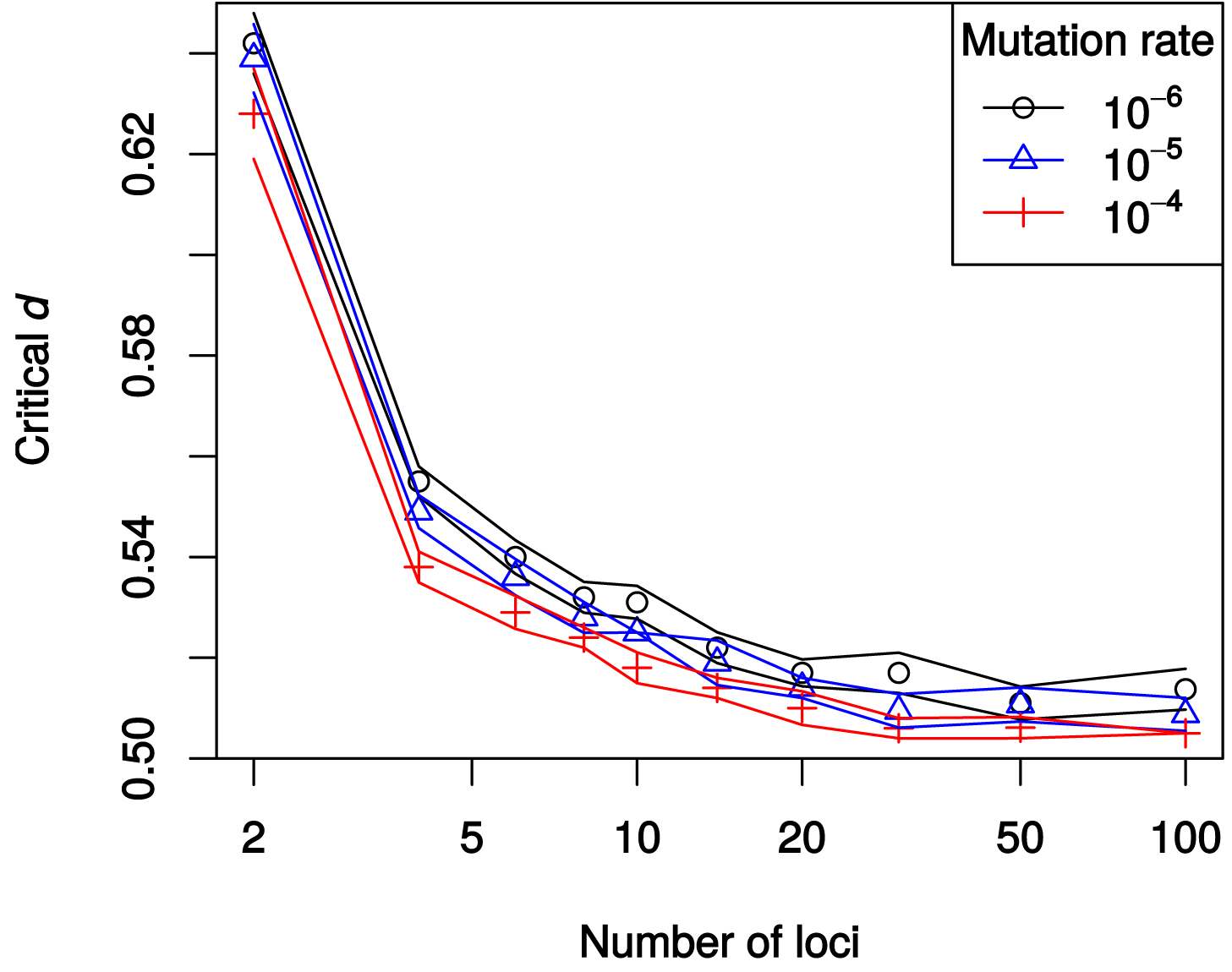
Critical value of the dominance parameter, *d_crit_*, such that the effective strength of balancing selection (*b_e_* in Eq. (14) in Methods), is positive (stable polymorphism) if *d > d_crit_*, and negative (unstable polymorphism) if *d* < *d_crit_*. Symbols represent means across replicates and lines represent averages ± two standard errors. Since the pattern for odd numbers of loci is more complex (see Fig. 6), only even values for the number of loci are included here. *N* = 1000, *y* = 2, *g* = 15.

From now on, we will focus on scenarios with large numbers of loci. For the case of 100 loci, Fig. 8 shows example allele-frequency trajectories with three different dominance parameters, *d*. For small *d*, each locus is almost fixed either for the summer or winter allele. For large *d*, all loci fluctuate at intermediate frequency. The critical dominance parameter, *d_crit_*, with 100 loci is close to 0.5, independently of the exponent of the fitness function (Fig. 9 A). For *d* < 0.5, i.e. if the currently favored allele is recessive, the effective strength of balancing selection is negative and polymorphism is unstable (Fig. 9 A). As *d* increases beyond 0.5, i.e. as the currently favored allele becomes more dominant, effective balancing selection becomes stronger (Fig. 9 A). Both the stabilizing and destabilizing effects increase with increasing exponent of the fitness function, *y* (Fig. 9 A). The finding of stable multi-locus polymorphism for *d* > *d_crit_* ≈ 0.5 is robust both to an imbalance in the number of generations in summer and winter (Fig. S5) and to seasonal changes in population size (Fig. S6).

**Figure 8:**
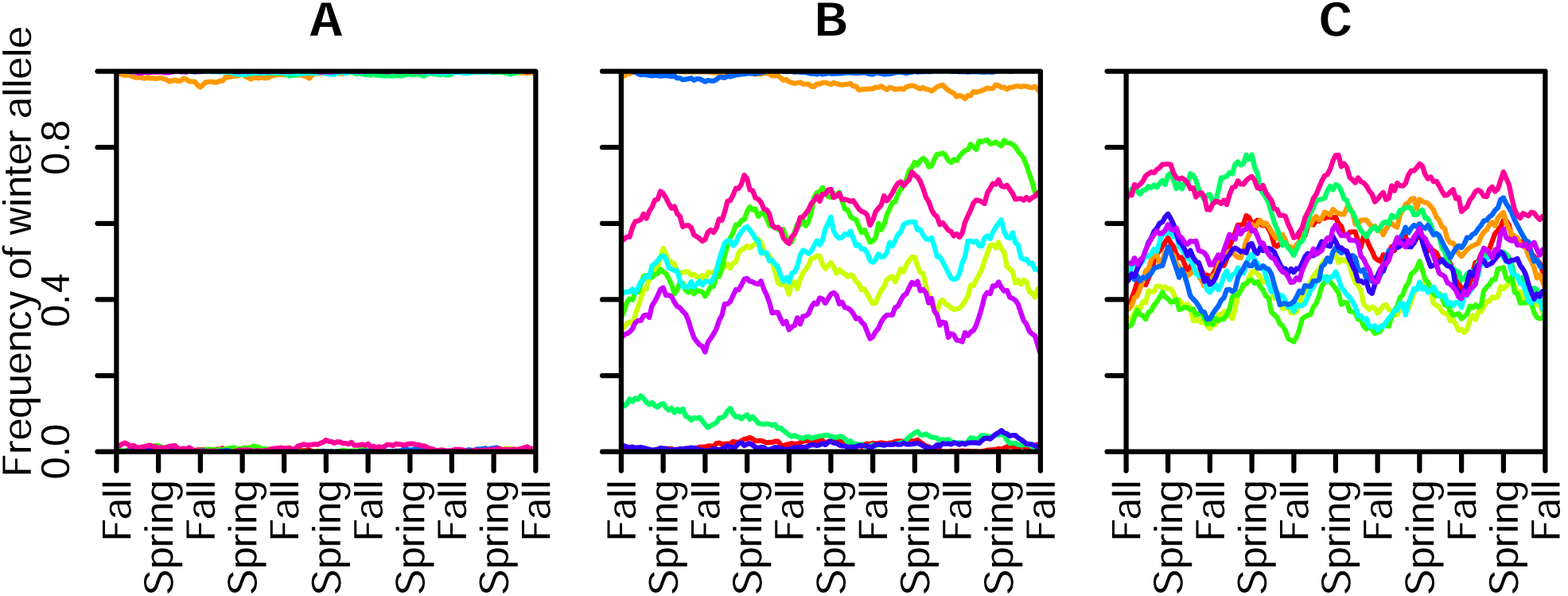
Three examples of allele-frequency trajectories for *N* = 1000, *L* = 100, *g* = 15, *y* = 4, *μ* = 10^−4^, and (A) *d* = 0.15, (B) *d* = 0.5, (C) *d* = 0.65. Only 10 randomly selected loci (shown in different colors) out of 100 loci are shown for five years (150 generations) in the middle of the simulation run (years 301 to 305).

**Figure 9:**
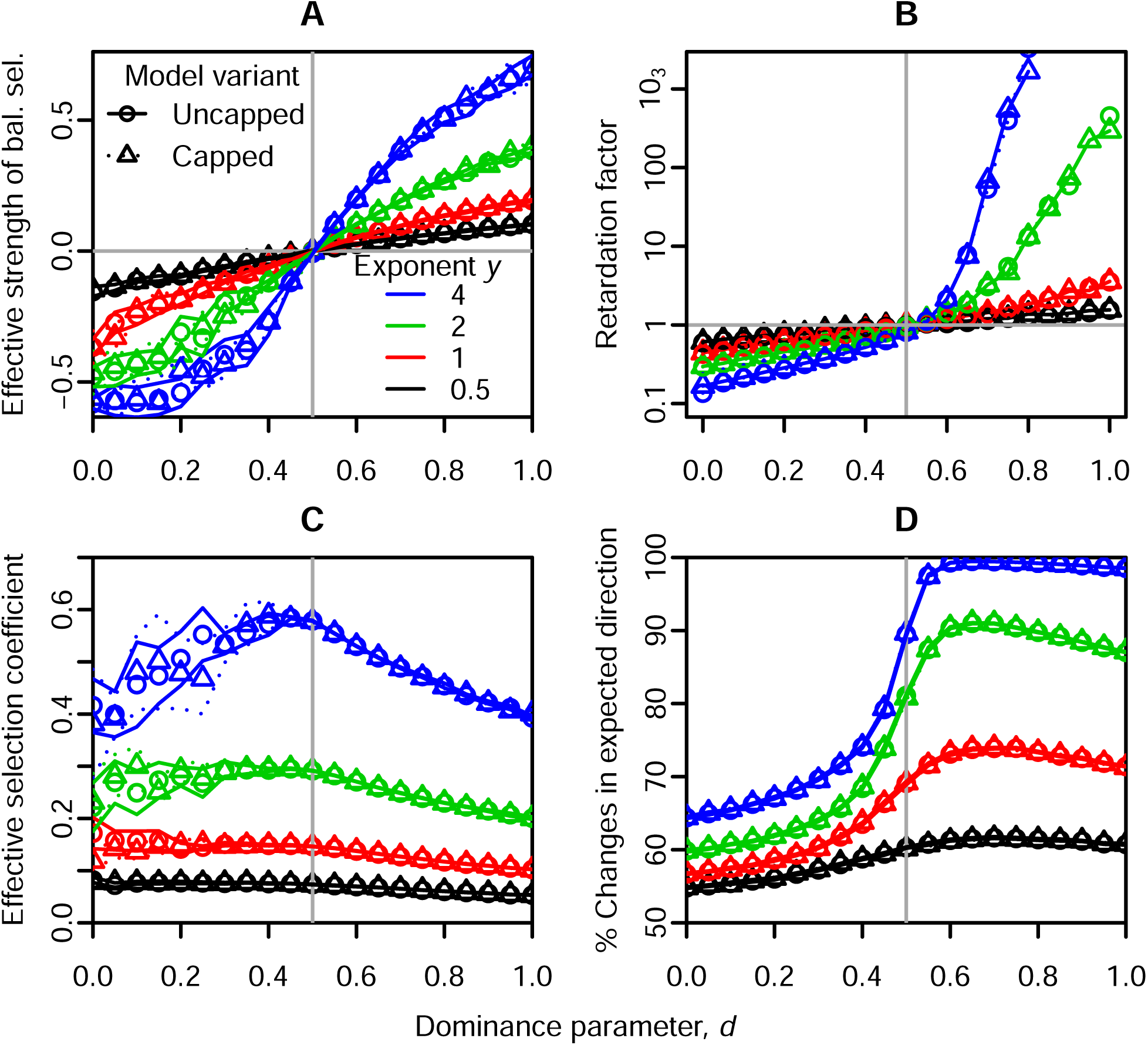
Influence of the dominance parameter *d* on (A) effective strength of balancing selection (*b_e_*, Eq. (14), Methods), (B) retardation factor, (C) magnitude of fluctuations (*s_e_*, Eq. (15), Methods), (D) predictability of fluctuations. Symbols indicate averages across replicates for the uncapped vs. capped model variant (often overlapping) and solid vs. dashed lines in A, C, and D indicate the respective means ± two standard errors. Lines in B simply connect maximum-likelihood estimates obtained jointly from all replicates. *N* = 1000, *L* = 100, *g* = 15, *μ* = 10^−4^. See Fig. S4 for more detailed information on the distribution and frequency-dependence of seasonal allele frequency changes. The vertical grey lines are at *d* = 0.5

A tendency for rare alleles to increase in frequency does not guarantee that the average lifetime of a polymorphism is larger than under neutrality [43, 47]. This is particularly interesting for fluctuating selection regimes with positive autocorrelation where alleles regularly go through periods of low frequency [43]. We therefore compute the so-called retardation factor [47], the average lifetime of a polymorphism in the selection scenario relative to the average lifetime under neutrality (see Appendix S2.3 for detailed methodology). The results for 100 loci are consistent with those for the effective strength of balancing selection: For *d >* 0.5, polymorphism under segregation lift is lost more slowly than under neutrality (Fig. 9 B).

To quantify seasonal fluctuations, we compute an effective selection coefficient (see Methods and Appendix S2.2). We also compute the predictability of fluctuations as the proportion of seasons over which the allele frequency changes in the expected direction, e.g. where the summer-favored allele increases over a summer season. Both the effective selection coefficient and the predictability of fluctuations have a maximum at intermediate values of *d* and increase with increasing exponent *y* of the fitness function Eq. (4) (Fig. 9 C, D). Also, effective strength of balancing selection, effective selection coefficient, and predictability of fluctuations increase with the number of generations per season (Fig. S7). Despite considerable variance in fitness across individuals in a population (coefficients of variation up to at least 0.25, see Fig. S8), the results for the capped model generally match the results for the uncapped model in all respects, especially for *d* > 0.5 (Fig. 9 A-D).

With increasing number of loci and with *d* > 0.5, the strength of balancing selection, the retardation factor, the magnitude and predictability of fluctuations all decrease (Fig. 10). Population size hardly influences effective strength of balancing selection and effective selection coefficient (Fig. 10 A, C), but large populations maintain polymorphism for longer (Fig. 10 B) and have more predictable allele-frequency fluctuations (Fig. 10 D). In small populations, polymorphism can even be lost slightly faster than under neutrality (Fig. 10 B).

**Figure 10:**
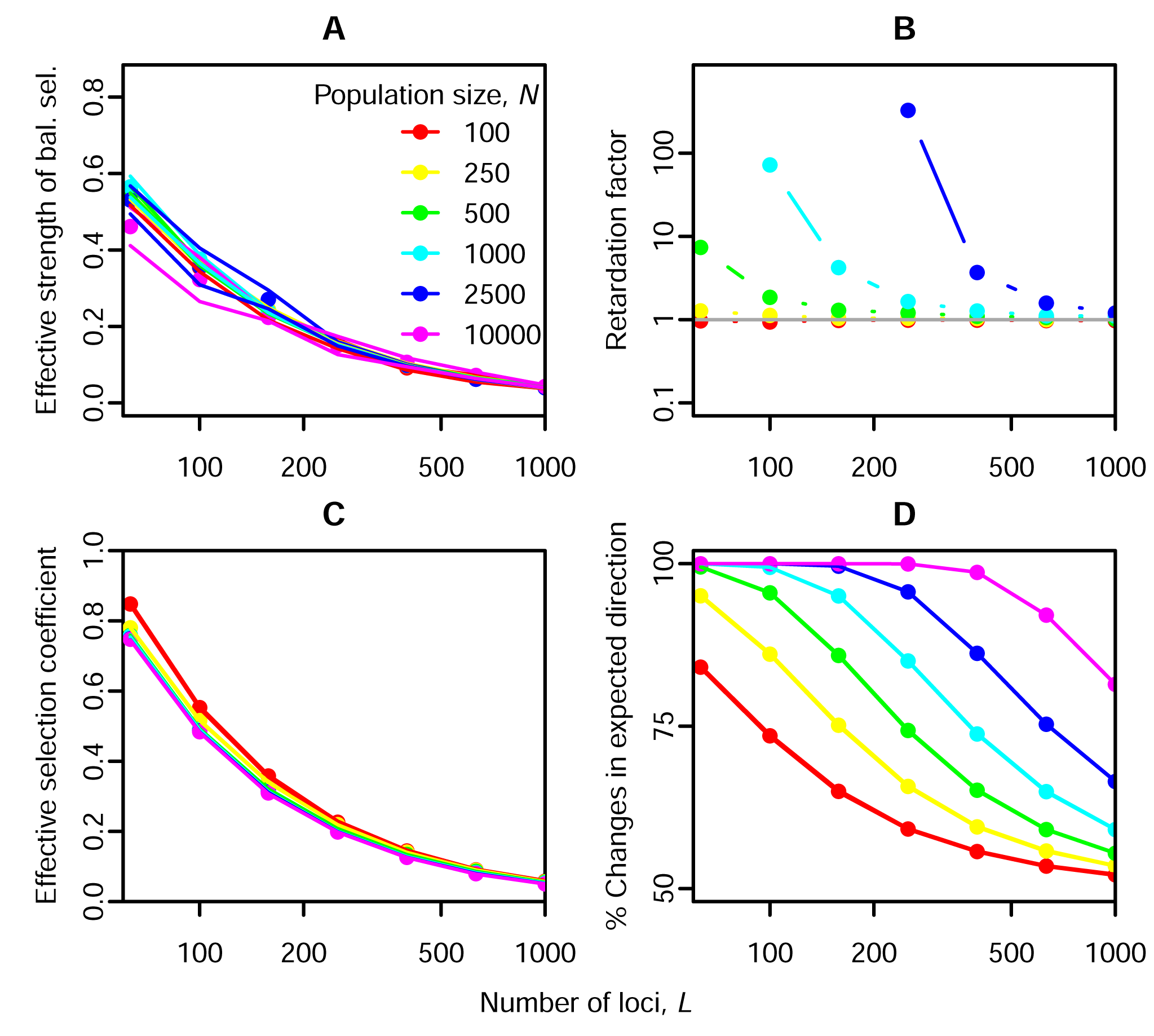
Influence of population size and the number of seasonally selected loci on (A) effective strength of balancing selection (*b_e_* in Eq. (14), Methods), (B) retardation factor, (C) magnitude of fluctuations (*s_e_* in Eq. (15), Methods), and (D) predictability of fluctuations. Symbols indicate averages across replicates and lines in A, C and D indicate means ± two standard errors (In C and D, standard errors are too small to be visible). Lines in B simply connect maximum-likelihood estimates obtained jointly from all replicates. Note that in (B) some points are missing because the rate of loss of polymorphism was too small to be quantified. *d* = 0.7, *y* = *4,g* = 15, *μ* = 10^−4^.

Finally, we consider a generalized model where parameters vary across loci and seasons. Independently for each locus *l*, we draw four parameters: summer effect size Δ_*s,ι*_ and winter effect size Δ_*w,l*_ are drawn from a log-normal distribution. Specifically, we draw the logarithms of these parameters from a bivariate normal distribution with mean 0, standard deviation 1, and correlation coefficient 0.9. Summer and winter dominance parameters, *d_s,l_* and *d_w,l_*, are drawn independently from a uniform distribution on [0,1]. Seasonal scores are then computed as 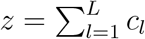, where the contribution *c_l_* of locus *l* in summer is 0 for winter-winter homozygotes, *d_s_*,_*l*_Δ_*s,l*_ for heterozygotes, and Δ_*s,l*_ for summer-summer homozygotes. Winter contributions are computed analogously. Because all effect sizes Δ are positive, all loci exhibit a trade-off between summer and winter effects. We use *y* = 4 here because it led to the most stable polymorphism in the basic model.

The results indicate that polymorphisms with different parameters can be maintained in the same population, with their allele frequencies fluctuating on various trajectories (Fig. 11 A). With a sufficiently high total number of loci, hundreds of stable polymorphisms (positive expected frequency change of a rare allele, see Appendix S2.4 for details) can be maintained in populations of biologically plausible size (Fig. 11 B). The number of loci classified as stable depends only weakly on population size. However, only a small proportion of the polymorphisms classified as stable also exhibit detectable allele-frequency fluctuations, defined as changes in the expected direction by at least 5 % in at least half of the seasons (Fig. 11 C,D). The number of detectable polymorphisms is highest at an intermediate total number of loci and increases with population size (Fig. 11 D). Detectable polymorphisms tend to have larger summer and winter effect sizes than polymorphisms that are only stable (Fig. 11 E). Compared to unstable polymorphisms, stable polymorphisms are more balanced in their summer and winter effect sizes (two-sample *t*-test on | In(Δ_*s,l*_/Δ_*w,l*_)|, *p <* 2.2 10^−16^, Fig. 11 E, see also Fig. S9). For almost all stable polymorphisms, detectable or not, the average dominance parameter across seasons is larger than 0.5 (Fig. 11 F).

**Figure 11:**
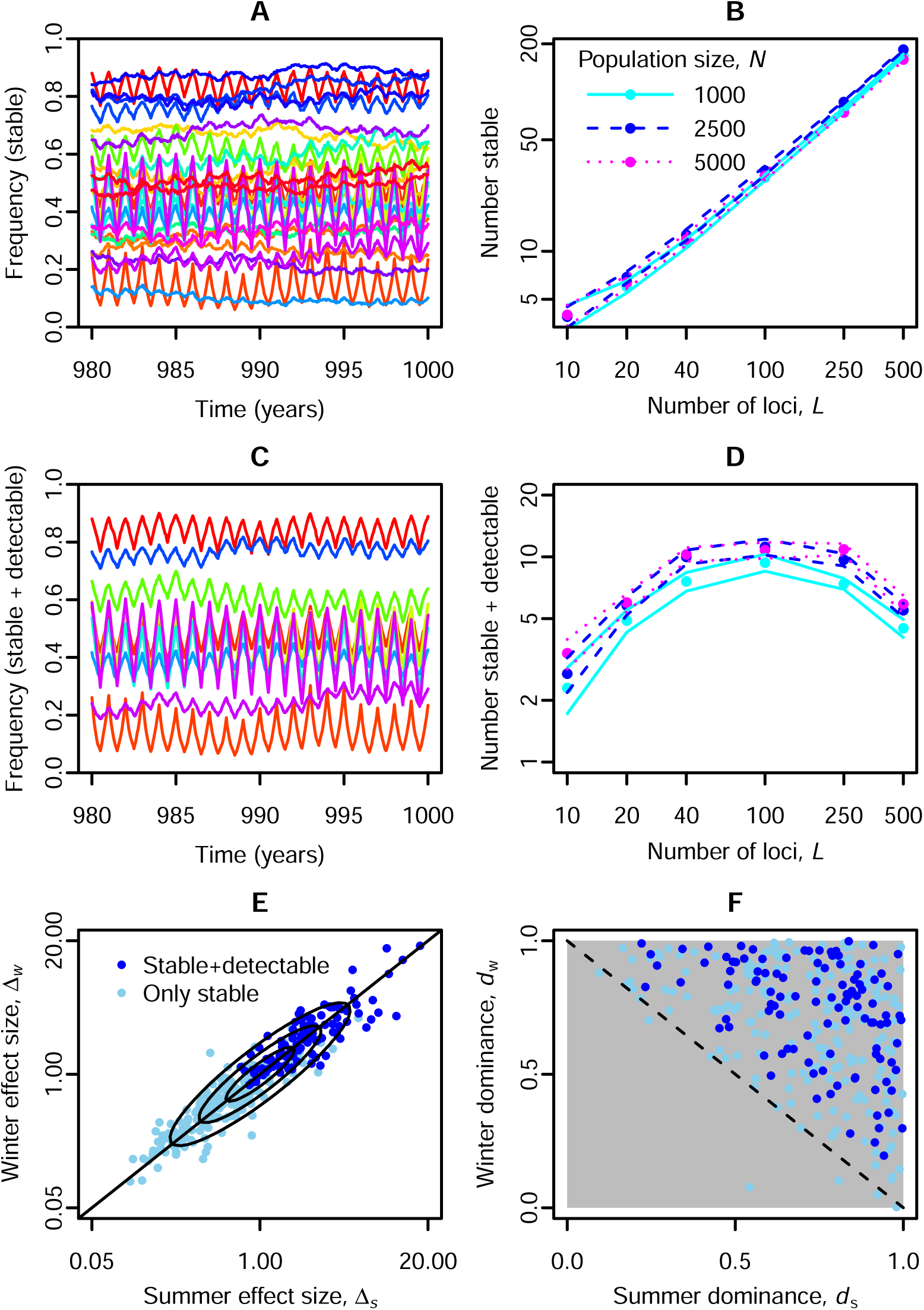
Stability of polymorphism and detectability of allele-frequency fluctuations when parameters vary across loci and seasons. A) Snapshot of allele-frequency trajectories for stable polymorphisms in one simulation run. B) Average number of stable polymorphisms as a function of the total number of loci for different population sizes. C,D) As in A and B, but only for polymorphisms that are also detectable. E) Winter effect size, Δ_*l,w*_, vs. summer effect size, Δ_*l,s*_, for stable and detectable and only stable polymorphisms. The plot shows pooled results over ten simulation runs with independently drawn parameters. Oval isoclines indicate the shape of the original sampling distribution, with 75 % of the sampling probability mass inside the outermost isocline. F) Corresponding dominance parameters. The dominance parameters were originally drawn from a uniform distribution on the unit square. Parameters: *y* = 4, *g* = 10, *μ* = 10^−4^ and in (A,C,E,F) *N* = 10, 000, *L* = 100.

## Discussion

We have studied a simple model for seasonally fluctuating selection that maps the multi-locus genotype to fitness in two steps. First, we count the number of loci homozygous for the currently favored allele and add the number of heterozygous loci weighted by a dominance parameter. The resulting seasonal score is then mapped to fitness via a monotonically increasing function which accounts for strength of selection and epistasis. The previously studied cases of multiplicative selection and selection on a fully additive phenotype are special cases of our model. We identified a general mechanism, which we call segregation lift, by which seasonally fluctuating selection can maintain polymorphism at tens or hundreds of unlinked loci. Segregation lift requires that the average dominance parameter of the currently favored allele, the summer allele in summer and the winter allele in winter, is sufficiently large. Individuals with many heterozygous loci then have higher average scores in both seasons than individuals with the same number of summer- and winter-favored alleles but more homozygous loci. Unlike in previously studied fully additive models, fully homozygous types thus cannot fix in the population and multilocus polymorphism is maintained.

The critical value of the dominance parameter required to maintain polymorphism, *d_crit_*, depends mostly on the type of epistasis and on the number of loci. Without epistasis, i.e. for multiplicative selection, *d_crit_* is 0.5. With synergistic epistasis it is close to one when there are multiple loci. For diminishing-returns epistasis and a small number of loci, *d_crit_* can be substantially above 0.5, but as the number of loci increases, *d_crit_* decreases toward 0.5. For one hundred loci, for example, only small deviations from additivity are required to maintain polymorphism at all loci.

## Robustness and plausibility of segregation lift as a mechanism to maintain variation

The conditions for stable polymorphism under segregation lift are insensitive to the mutation rate (see Fig. 7) and to the number of generations per season, even if there is an asymmetry (see Figs. S7 and S5). Segregation lift is also robust to seasonal changes in population size (Fig. S6) and to variation in effect sizes and dominance parameters across loci (Fig. 11). Future work will be needed to assess how linkage between selected loci affects stability of polymorphism under segregation lift.

Whenever there is balancing selection at a large number of loci, genetic load is a potential concern. In the case of segregation lift with diminishing-returns epistasis, however, genetic load does not appear to play an important role. The results for our capped model closely match the results for the original, uncapped model. Thus, although individuals might occasionally have a large number of offspring in the uncapped model, large offspring numbers are not required for stable multi-locus polymorphism or any of the other observed features.

Although changes in dominance across seasons or traits, a key requirement for segregation lift, have traditionally been considered rare [42, 48], the required changes are small and we argue that they can indeed arise naturally in situations with antagonistic pleiotropy and seasonal changes in the relative importance of traits (see Fig. 3). In reality, it is likely that seasonal differences are more pronounced in some years than in others, and there may also be spatial variation in seasonality. If seasonality is weak and both traits are roughly equally important throughout the year, antagonistic pleiotropy can even lead to simple heterozygote advantage [Fig. S1 B, 41]. Thus, maintenance of polymorphism by segregation lift should also be robust to variation in the degree of seasonality. Segregation lift can also be interpreted as a specific type of phenotypic plasticity, where the ability of a genotype to adjust plastically to both summer and winter environments increases with the number of heterozygous loci. This plasticity could be mediated by environment-dependent changes in dominance for gene expression as observed in *Drosophila* [45].

All in all, the notion that seasonally fluctuating selection cannot contribute substantially to genomic variation appears no longer justified. The conditions for polymorphism in our model are much less restrictive than in previously studied scenarios. We do not need multiplicative selection but can allow other types of epistasis and the required small changes in dominance can arise naturally under antagonistic pleiotropy. Segregation lift is not affected by problems of genetic load and is robust to many perturbations in the parameters across time and loci. Because of this robustness, segregation lift should be able to maintain polymorphism at many loci in the genome under seasonally fluctuating selection and potentially also under other forms of temporal heterogeneity.

## Magnitude and detectability of allele-frequency fluctuations

Segregation lift not only explains stability of polymorphism, but can also produce strong and predictable seasonal fluctuations in allele frequencies. This is the case especially for large exponents *y* of the fitness function (Eq. (4)), our measure for the strength of selection, and for dominance parameters, *d*, slightly above 0.5. For even higher values of *d*, fluctuations are not as strong, presumably because heterozygotes are fitter and therefore more copies of the currently disfavored allele enter the next generation.

The magnitude of allele-frequency fluctuations also decreases with the number of loci under selection. This explains why the number of detectable polymorphisms is maximized at an intermediate number of loci in Fig. 11. In Appendix S3, we use a heuristic mathematical argument to explore the relationship between number of loci and magnitude of fluctuations in a population of infinite size. We show that as the number of loci goes to infinity, the effective strength of selection at each locus is expected to go to zero, i.e. effective neutrality. This is because more loci lead to higher overall seasonal scores, *z*, which under diminishing-returns epistasis leads to weaker average selection pressures at each locus. These findings and also our results in Fig. 11 suggest that even if segregation lift contributes substantially to maintaining polymorphism at a large number of loci, it is not necessarily easy to detect individual selected loci based on their allele-frequency fluctuations.

## Empirical evidence, alternative hypotheses, and potential tests for segregation lift

Recently, Bergland et al. [37] observed strong and stable allele-frequency fluctuations at a large number of loci in a temperate population of *Drosophila melanogaster.* The population was sampled twice a year for three successive years, once in spring and once in fall. Pooled whole-genome sequencing revealed hundreds of apparently seasonally selected single-nucleotide polymorphisms (SNPs). The SNPs were distributed throughout the genome, and most were not closely linked. At each SNP, one allele was favored during summer and the other allele during winter with large allele-frequency fluctuations (~10% over a single season of about 10 generations). As mentioned above, many of the SNPs are shared with African populations of *D. melanogaster* and some even with the closely related *Drosophila simulans,* suggesting long-term stability. Based on our results, it is plausible that seasonally fluctuating selection with segregation lift, even in the absence of other stabilizing mechanisms, can maintain polymorphism at such a large number of loci. It is less clear, however, whether segregation lift can explain the observed large magnitude of allele-frequency fluctuations. Only few loci exhibited such large fluctuations in the generalized model with parameter variation across loci (see Fig. 11 C, D). However, in our basic model with equal parameters across loci we did observe 100 loci with allele-frequency fluctuations of similar magnitude as in the *Drosophila* data (see Fig. 8 C), suggesting that, in principle, segregation lift is compatible with large fluctuations at many loci.

Of course, segregation lift is not the only possible explanation for polymorphisms with seasonal fluctuations in frequency. Such fluctuations are also possible under stabilizing selection on an additive trait with a temporally fluctuating optimum because recurrent mutation can induce substantial genetic variance [32, 49]. However, this is really a form of mutation-selection balance and not a mechanism leading to long-term stable polymorphism. Alternative mechanisms that could potentially lead to both fluctuations and long-term stability are 1) differential responses to fluctuating resource concentrations and population densities [19, 50–52], 2) a so-called “temporal storage effect” where genetic variation can be buffered by a long-lasting life-history stage on which selection does not act, or by some other protected state [51, 53, 54]. However, these mechanisms are more commonly studied in an ecological context, and it is unclear whether they can maintain polymorphism at multiple loci.

Detecting segregation lift in empirical data will be challenging, but should be possible in principle. One way would be to stock outdoor cages with various multi-locus genotypes and track fitness over one or more seasons. The main challenge is that the pivotal dominance parameter, *d*, is relative to the seasonal score, *z*, which mediates between multi-locus genotype and fitness, and is itself not directly measurable. Since the shape of the fitness function, *w*, is also generally unknown, it is not possible to infer *d* from fitness measurements of different single-locus genotypes in a common genetic background (see Fig. 4). For a set of loci with the same effect sizes and dominance parameters, we would be able to compare multi-locus genotypes with the same number of summer and winter alleles and conclude that there is segregation lift if fitness in both seasons increases with the number of heterozygous loci. More generally, given fitness measurements for a large number of multi-locus genotypes, we could use statistical methods such as machine learning to jointly estimate parameters of the fitness function, effect sizes, and dominance parameters and thereby assess whether or not there is segregation lift. Such statistical approaches could also take into account the existence of several multiplicative fitness components each with a set of contributing loci that might exhibit segregation lift and epistatic interactions. In any case, one point is clear: Looking at one locus or two loci at a time is not sufficient to understand the multi-locus phenomenon of segregation lift.

## Conclusions

We have identified segregation lift as a general mechanism by which seasonally fluctuating selection can maintain polymorphism at hundreds of unlinked loci in populations of biologically reasonable size. Segregation lift circumvents the problems associated with maintenance of polymorphism under stabilizing selection and does not require highly heterozygous individuals to have unrealistically many offspring. Given the ubiquity of environmental fluctuations, this mechanism could make a substantial contribution to genetic variation in natural populations of many taxa. With easy access to population genomic data across time and space, we are now in a good position to make progress on the puzzle of genetic variation, that is quantify the contributions of segregation lift and other mechanisms to overall genetic variation.

## Methods

For the basic model and the capped model, we assess stability of polymorphism by estimating an effective strength of balancing selection, *b_e_*, from the year-to-year allele-frequency dynamics. For this, we fit a standard balancing selection model [55] of the form

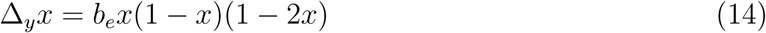

to average changes in allele frequency over one yearly cycle, Δ_*y_x_*_ (see Appendix S2.2 for details). Positive values of *b_e_* indicate that rare alleles tend to become more common in the long run, whereas negative values indicate that rare alleles tend to become even more rare. Second, we quantify the magnitude of fluctuations over individual seasons. For this, we fit a standard directional selection model [55]

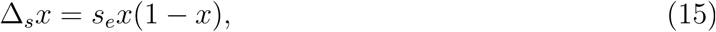

with an effective selection coefficient *s_e_,* to average allele-frequency changes over one season, Δ_*S*_*x* (see Appendix S2.2).

To obtain a measure for statistical uncertainty in our results, we run 10 replicates for every parameter combination and calculate effective strength of balancing selection, *b_e_*, and effective selection coefficient, *s_e_*, independently for each replicate. We do the same for the predictability of fluctuations, i.e. the proportion of seasons in which a locus changes its allele frequency in the expected direction. In all three cases we report the mean over replicates ± two standard errors of the mean. To obtain retardation factors, we run 100 replicates until the first polymorphism is lost or a maximum time of 500 years is reached. From the times of loss for the replicates, we obtain maximum likelihood estimators for the rate of loss of polymorphism (see Appendix S2.2 for details).

C++ source code and R scripts for the individual-based simulations and their analysis are available upon request.

## Acknowledgments

MJW gratefully acknowledges fellowships from the Stanford Center for Computational Evolutionary and Human Genomics (CEHG) and from the Austrian Science Fund (FWF). For helpful discussion and/or comments on the manuscript, we would like to thank Michael Desai, Joachim Hermisson, Oren Kolodny, Mike McLaren, Pleuni Pennings, Jitka Polechová, and members of the Petrov lab. Simulations were performed on Stanford’s FarmShare Cluster and on the Vienna Scientific Cluster.

## Appendix S1 Local stability analysis for the additive scenario (*d* = 0.5) with diminishing-returns epistasis

In this section, we consider a scenario where alleles contribute to the seasonal score *z* additively within and between loci (*d_s_* = *d_w_* = 0.5 in Eq. (1) and Eq. (2)). We show that in the deterministic model without recurrent mutation, a population fixed for a balanced haplotype with *L/2* summer-favored alleles and *L/2* winter-favored alleles (*L* even) cannot be invaded by any other haplotype. We give an intuitive rather than a mathematically formal outline of the stability analysis.

Assume that a certain balanced haplotype is at frequency close to 1 in the population. We will call it the resident haplotype. Let *ϵ*_*i,j*_ be the combined frequency of all haplotypes that have a winter allele at *i* positions where the resident haplotype has a summer allele and a summer allele at j positions where the resident haplotype has a winter allele. Thus an *i,j*-haplotype has *L*/2 + *i* – *j* winter alleles, *L*/2 – *i* + *j* summer alleles, and differs in *i* + *j* positions from the resident haplotype.

Rare *i*, *j*-haplotypes will almost certainly occur in a genotype with a copy of the resident balanced haplotype. This has two consequences. First, recombination with the resident balanced haplotype will produce other haplotypes that differ in at most *i* + *j* positions from the resident haplotype. Haplotypes with more than *i* + *j* differences cannot be produced because if resident and invading haplotype have the same allele at a locus, then all offspring haplotypes will carry this allele as well. Given free recombination, the probability that one of the two recombinant haplotypes produced by an *i*, *j*-resident genotype (consisting of one *i*, *j*-haplotype and one resident haplotype) is of type *i*, *j* is 1/2^*i*+*j*^. Second, the relevant fitness values for the invasion of the *i*, *j*-haplotype are *w*((*L* + *i* – *j*)/2) in winter and *w*((*L – i* + *j*)/2) in summer. The population mean fitness will be approximately *w*(*L*/2) in both seasons, namely the fitness of the resident-resident genotype.

To linear order, the frequency of the *i*, *j*-haplotype after one yearly cycle with *g* generations of summer and g generations of winter will be

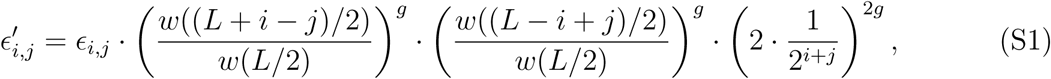

where the factor of two in the last term comes from the fact that each genotype contributes on average two haplotypes in the next generation. From the assumption that intermediate types are favored in the long run (negative epistasis), we can conclude that

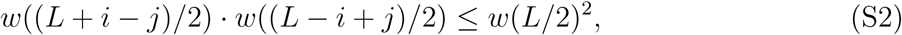

with equality for *i* = *j*. Hence 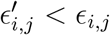 for *i* ≠ *j*, showing that unbalanced haplotypes cannot invade the population.

For a balanced haplotype (*i* = *j*),

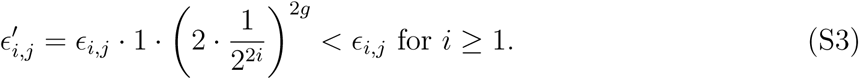

Intuitively, other balanced haplotypes cannot invade, because they differ from the resident balanced haplotype at more than one position and are therefore broken down by recombination. Hence we have shown that a population fixed for a certain balanced haplotype cannot be invaded by any other balanced or unbalanced haplotype.

**Figure S1:**
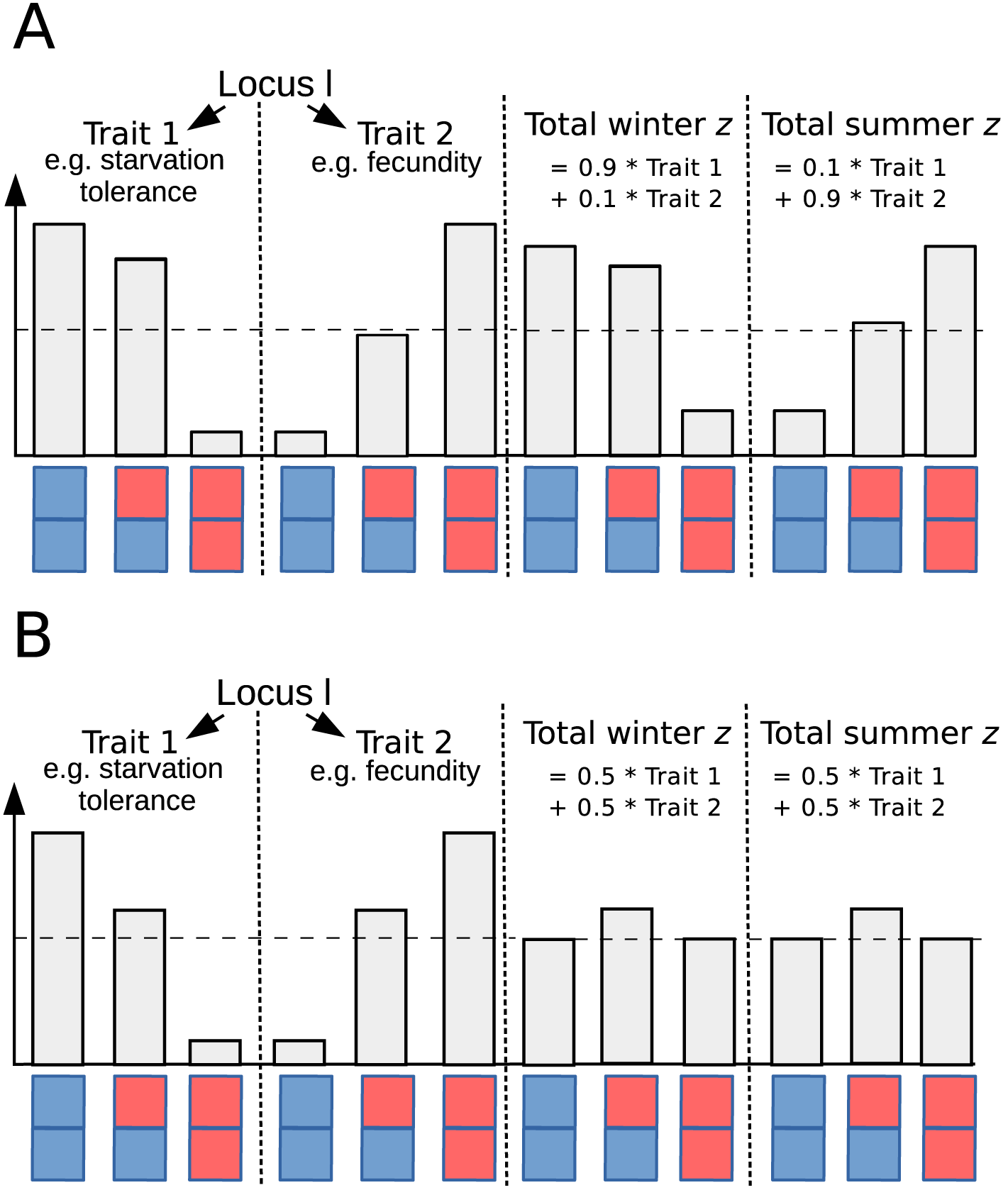
Alternatives to the scenario in Fig. 3. A) If heterozygotes are relatively close to the fitter homozygote with respect to one of the traits and the beneficial allele is only slightly recessive for the other trait, we can still obtain a beneficial reversal of dominance with respect to the seasonal score, z. However, in this case, *d_s_* ≠ *d_w_*. B) If the two traits are of similar importance in the two seasons, heterozygotes are fitter than either homozygote at all times.

## Appendix S2 Detailed methods

### Individual-based simulations

We assume discrete, non-overlapping generations with population size *N* and *L* loci. In each generation, the following events take place in this order: 1) The fitnesses of all individuals in the parent generation are calculated using the genotype-to-fitness map described in the main text (Eq. (1), Eq. (2), Eq. (4)). 2) Each individual in the offspring generation draws two parents, independently, with replacement (i.e. selfing is possible), and proportionally to parent fitness. 3) Each parent passes one set of alleles to each of its offspring. We assume unlinked loci such that at each of the *L* loci each of the two allele copies is passed on with equal probability, independently of the alleles passed on at other loci, and also independently of which alleles were passed on to other offspring of the same parent. 4) Independently and with probability *μ*, each of the allele copies passed to an offspring mutates to the respective other allele. 5) The parent generation is removed from the model and replaced by the individuals in the offspring generation.

To initialize a simulation run, we randomly assemble genotypes. Independently for each individual, locus, and allele copy, we draw the summer and the winter allele with equal probabilities. Each simulation runs for 500 years, corresponding to 500 2 *g* generations. Each parameter combination is replicated 10 times. For each locus, we store the allele-frequency trajectory over time.

### Stability of polymorphism and magnitude of fluctuations

To quantify and compare the dynamics of different simulation runs, we compute three summary statistics from the allele-frequency trajectories: an effective strength of balancing selection, *b_e_*, as a measure of the stability of polymorphism, an effective selection coefficient, *s_e_,* as a measure of the magnitude of fluctuations, and the proportion of allele frequency changes that go in the expected direction as a measure of predictability of fluctuations.

To compute the strength of balancing selection, we divide the allele-frequency interval between 0 and 1 into 25 equally-sized bins (from 0 to 0.04, from 0.04 to 0.08, …). For each frequency bin, we compute the average allele-frequency change over the course of one year, from the middle of the cold season to the middle of the next cold season. We chose the middle of the cold season as a reference point because at this point the average frequency across all loci should be approximately 0.5. Choosing the middle of the warm season as the reference point should give the same results, but choosing either the beginning or end of the warm season would lead to asymmetric results. We average over all loci and times, for which the frequency at the reference point is in the respective frequency bin. Because we want to quantify dynamics at equilibrium, we only use data from the second half of each simulation run. We also exclude data points for which the allele frequency at the beginning of the year is exactly 0 or 1. For each bin, we then subtract an approximate average frequency change due to mutations 2 · *g* · (*μ* · (1 – *p̅*) *– μ · p̅*) = 2 · *g · μ* · (1 – 2*p̅*), where *p̅* is the midpoint of the respective frequency bin (0.02, 0.06,…). Finally, we use the lm function in R [56] to fit a balancing-selection model of the form *b_e_ p̅* (1 – *p̅*)(1 – 2*̅*) to the mutation-adjusted average allele-frequency changes. The coefficient b_e_ is our effective strength of balancing selection. Example model fits are shown in Fig. S2 A.

**Figure S2:**
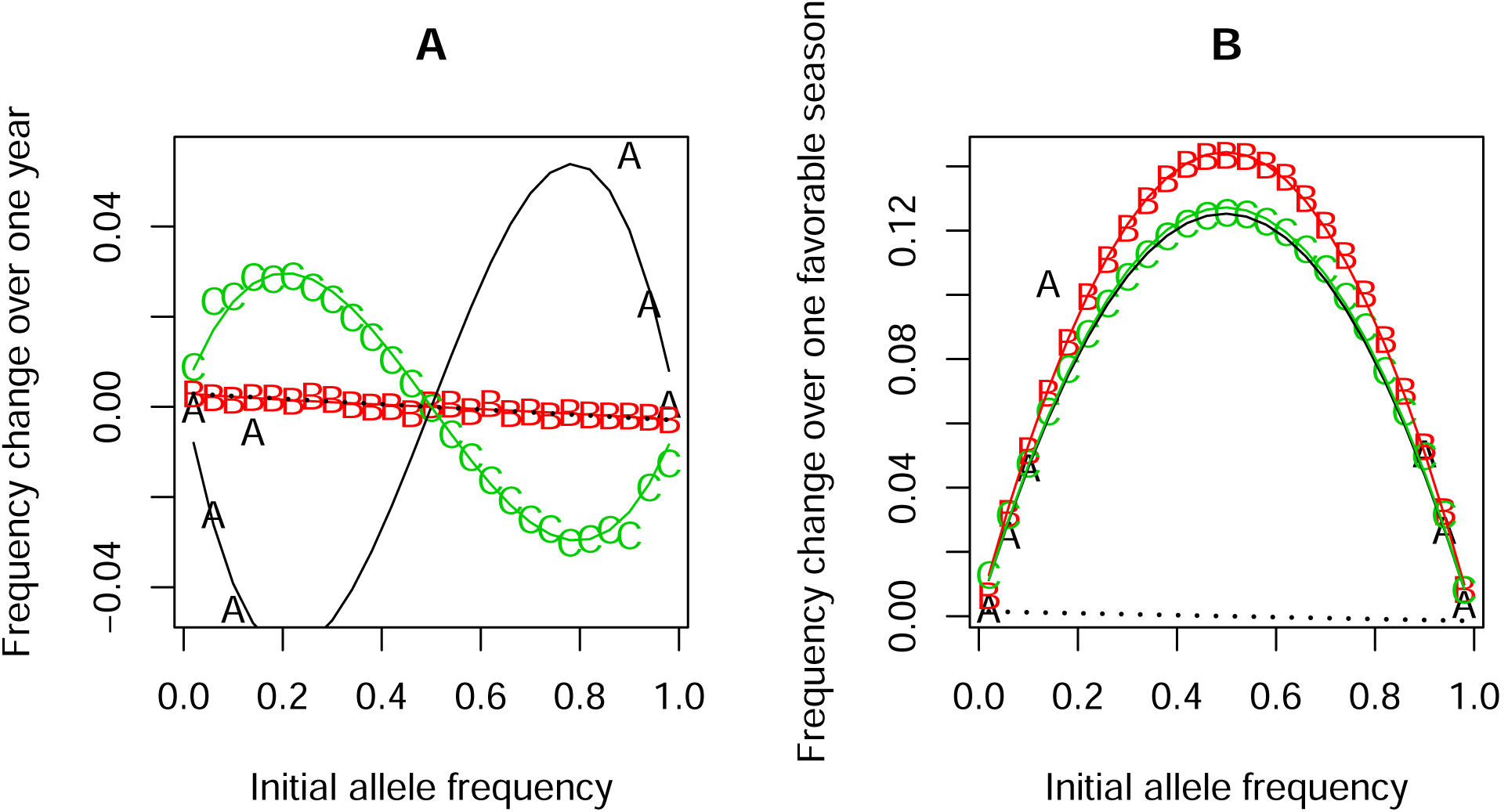
Examples of model fits to estimate (A) the effective strength of balancing selection (Eq. (14)) and (B) the effective selection coefficient (Eq. (15)). The letters A, B, and C in the plots indicate the panel in Fig. 8 that depicts the corresponding time series.

To compute the effective selection coefficient, we use the same data subsets and frequency bins as for the effective strength of balancing selection. But now we compute the average frequency change of the currently favored allele for each season, i.e. of the summer allele from spring to fall or of the winter allele from fall to spring. We bin data points according to the midseason frequency of the currently favored allele. After subtracting *g*·*μ*· (1 – 2*p̅*), the approximate expected frequency change due to mutations, we fit the model *s_e_·p̅* (1 – *p̅*). The coefficient *s_e_* is the effective selection coefficient. Example model fits are shown in Fig. S2 B.

### Retardation factor

To obtain the retardation factor, we ran additional simulations without recurrent mutations (*μ* = 0). We started at allele frequency 0.5 for all loci and simulated *n*_rep_ = 100 replicate populations. We stopped the simulation when polymorphism was lost at the first locus, but at most after 500 years, corresponding to *t*_max_ = 500·2· *g* generations. For each parameter combination, we ran a neutral control simulation, which was achieved by setting *y* = 0.

The relevant results for each parameter combination are the number of replicates, *n*_lost_, in which the first polymorphism was lost before *t*_max_ and the times of loss for each of these replicates *t*_1_,*t*_2_,…,*t_n_lost__*.

For simplicity, we assume that polymorphism is lost with the same probability *p* in every generation such that the time to the first loss is geometrically distributed. We then adopt a Maximum-Likelihood approach to estimate *p*. The likelihood of *p* is

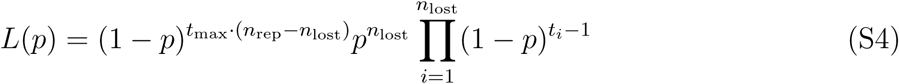

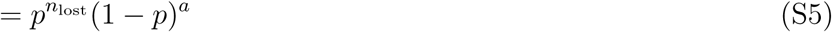

With

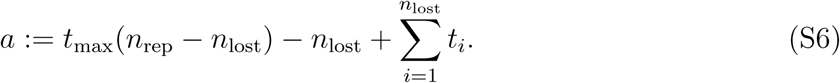

To look for extreme values, we take the first derivative with respect to *p*

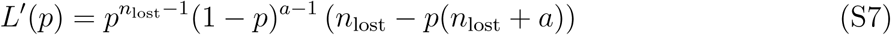

and obtain our estimator

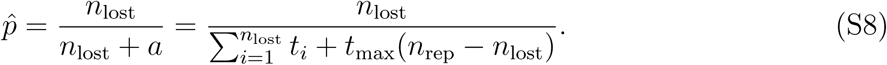

Eq. (S7) is positive for *p* < *p̂* and negative for *p* > *p̂.* Therefore, *p̂* is a maximum of the likelihood function. To see that this estimator makes sense, consider an example where loss is so fast that all replicates lose a polymorphism before *t*_max_. In this case, *p̂* will be the inverse of the average time to loss.

For each parameter combination, we obtain the ML-estimator *p̂*_sel_ under seasonally fluctuating selection and the ML-estimator for the corresponding neutral scenario *p̂*_neutral_. Finally, we compute the retardation factor as *p̂*_neutral_/*p̂*_sel_. Values larger than one occur for selection scenarios that tend to maintain polymorphism longer than neutrality, whereas in scenarios with a retardation factor below one polymorphism is lost more quickly than under neutrality.

### Stability in asymmetric scenarios

In our model with parameter variation across loci and seasons, and also in the basic model with different numbers of generations in summer and winter, the dynamics are generally asymmetric. That is, either the summer or the winter allele is more common on average. Hence the standard balancing selection model Eq. (14) does not fit anymore. We therefore use a different method to assess stability of polymorphism for individual loci. For each frequency bin between 0 and 0.5, we take all time points for which the frequency of the currently rare allele is in the frequency bin at the beginning of the cycle and compute the average change in its allele frequency over one cycle. As above, we subtract the expected input from new mutations. We do this separately for 10 replicate simulation runs (or for 10 disjoint subsets of the data if the actual replicates differ in the locus parameters) and for each frequency bin we compute the interval mean ± 2 standard errors. We then call a polymorphism stable if the rare allele has a positive expected frequency change in the most marginal frequency bin whose interval does not overlap zero.

## Appendix S3 Magnitude and detectability of seasonal fluctuations with diminishing-returns epistasis

In this section, we use a heuristic mathematical argument to explore how the magnitude of allele-frequency fluctuations changes as the number of loci increases in a population of infinite size.

Let us assume that there are *L* +1 loci in total, with one focal locus whose dynamics we will now study while the remaining *L* loci form its “genetic background”. Let us first consider the contribution, *z_b_,* of the genetic background to the seasonal score, *z,* i.e. the total score minus the contribution from the focal locus. For simplicity, we will focus on the expected allele frequency change over one generation at one of the two time points at which the allele frequency equals the yearly average allele frequency. At this point, both the population mean and the variance of the score *z_b_* among individuals in the population should be approximately proportional to the number of loci:

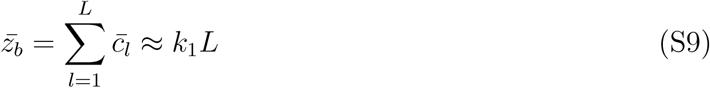

And

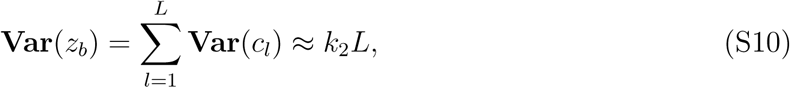

where we have assumed free recombination. Here, *c_l_* is the contribution of locus *l* to an individual’s score, and *k*_1_ and *k*_2_ are positive constants that will depend on the distribution of locus parameters. For example, if all loci have symmetric parameters (Δ_*s,l*_ = Δ_*w,l*_ =: Δ_*ι*_ and *d_s,l_* = *d_w,l_* =: *d_l_*), all loci will have an allele frequency of approximately 0.5 in the middle of summer. Since mating is random, allele frequencies will be in approximate Hardy-Weinberg equilibrium in every generation before selection and, with ~ denoting averages across loci,

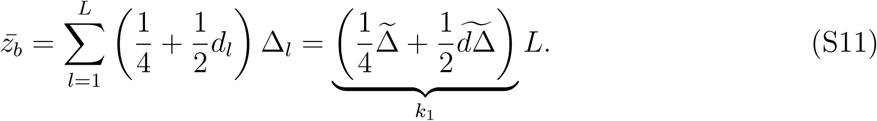

Similarly,

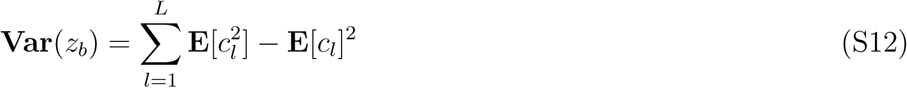

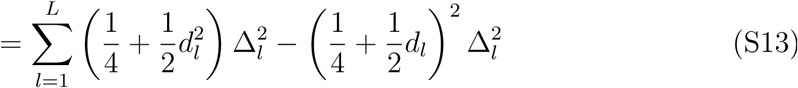

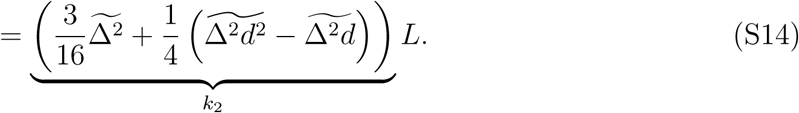

Next, we quantify the fitnesses of the three genotypes (WW, WS, SS) at a focal locus averaged over the possible genetic backgrounds. For this, we expand the fitness function *w*(*z*) as a second-order Taylor expansion around *z¯_b_*:

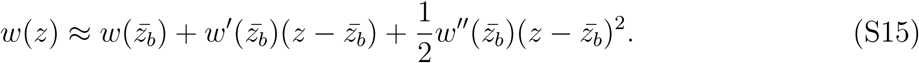

Now, we need the values of the phenotype *z* for the three genotypes at the focal locus. Let us assume that the focal locus has parameters *d* and Δ. Because we consider the population in the middle of summer, the winter-winter homozygote (WW) at the focal locus does not contribute anything to the seasonal score and *z* = *z_b_.* For the heterozygote (WS), *z* = *z_b_* + *d* Δ, and for the summer-summer homozygote, *z* = *z_b_* + Δ. With this, we obtain

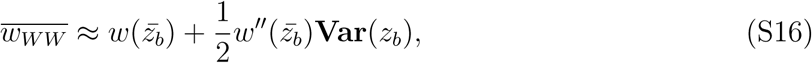

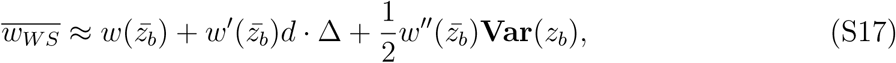

and

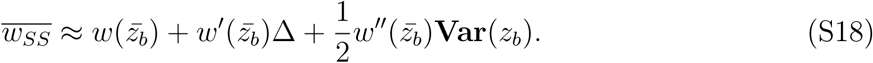

Finally, we compute the expected change in summer allele frequency, *p*, over one generation at the focal locus. The frequency of the summer allele in the next generation is and the proportional change in allele frequency is

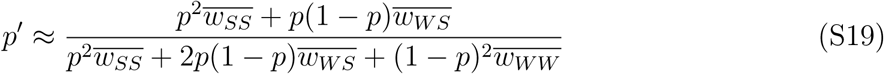

and the proportional change in allele frequency is

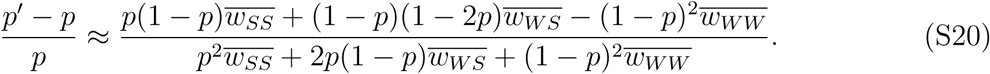

Substituting Eq. (S16)–Eq. (S18) and simplifying, we obtain

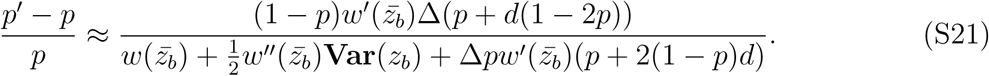

For the specific fitness function fitness function used in this paper

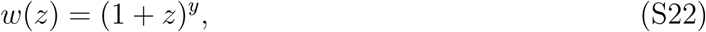

with

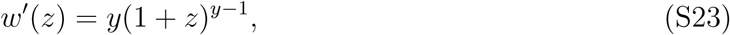

and

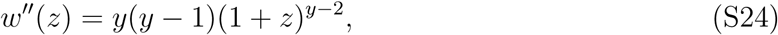

and using Eq. (S9) and Eq. (S10), we obtain

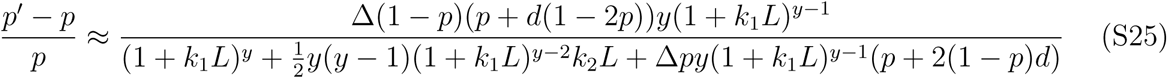

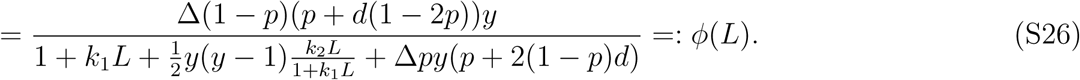

It follows that,

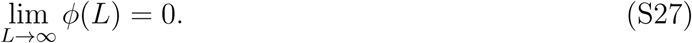

Moreover,

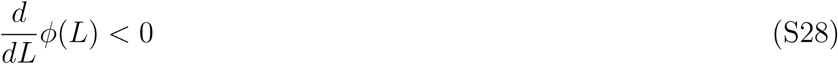

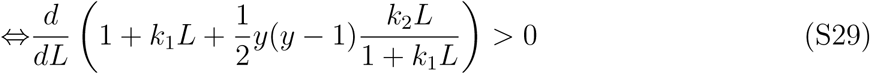

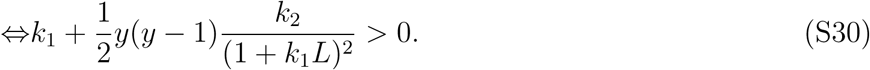

With *y >* 1, the second term is positive and the inequality is always fulfilled. With *y <* 1 it is fulfilled for sufficiently large *L*. Thus, with *y* > 1, the magnitude of allele-frequency change in an infinite population decreases monotonically as *L* increases and approaches zero for very large numbers of loci. With *y* < 1, allele-frequency change may first increase with increasing number of loci, but eventually decreases and also approaches zero. This finding indicates that there is a limit to the number of loci that can be detected to exhibit seasonal allele-frequency fluctuations. Intuitively, as the number of loci increases the average fitness of the genetic background becomes larger. Thus selection at a focal locus gets effectively weaker.

## Appendix S4 Additional results

See Figs. S3–S9.

**Figure S3:**
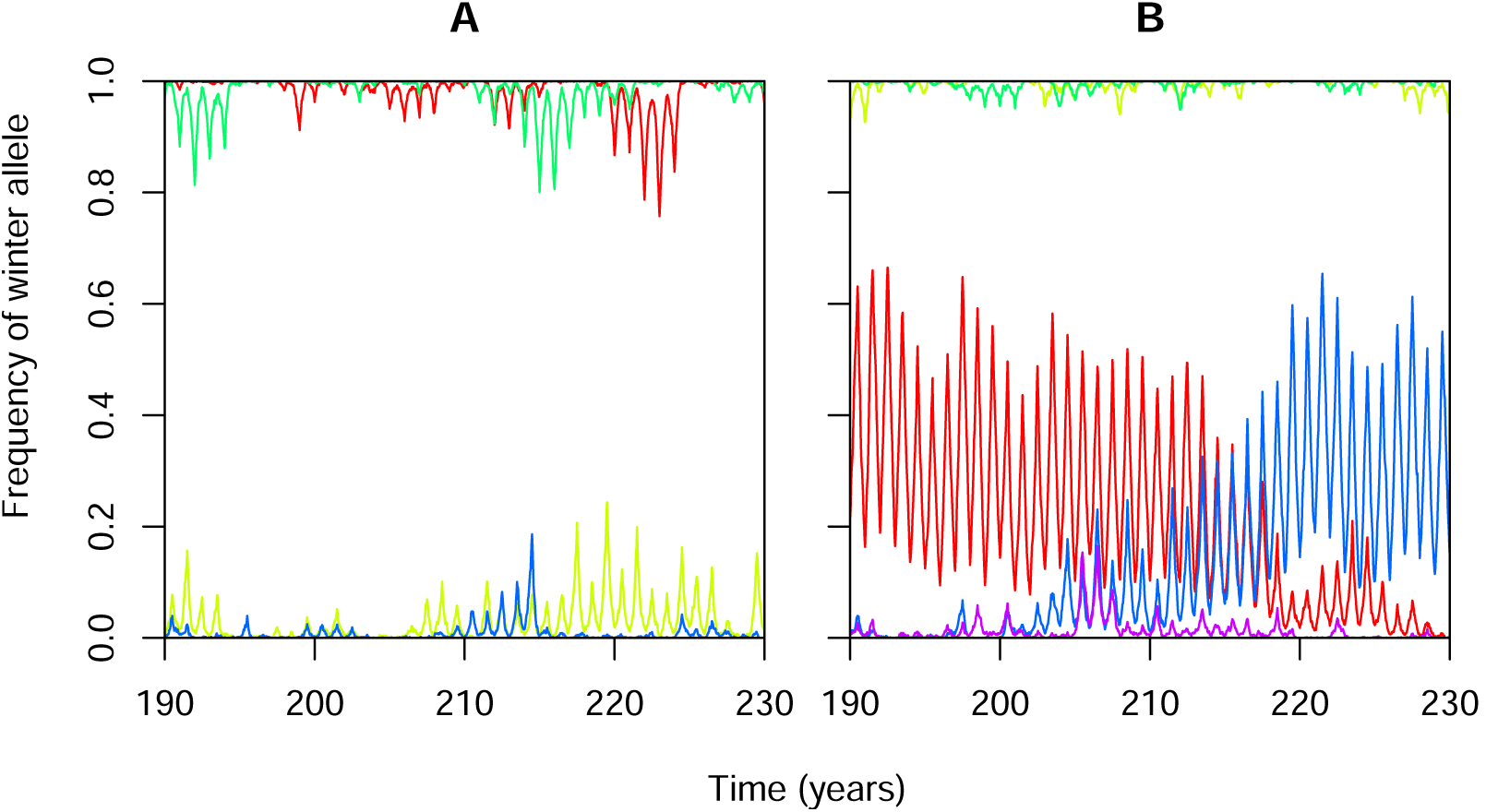
Example time series for the additive model (*d* = 0.5) with (A) four loci, or (B) five loci. *N* = 1000, *g* = 15, *y* = 1, *μ* = 10^−4^.

**Figure S4:**
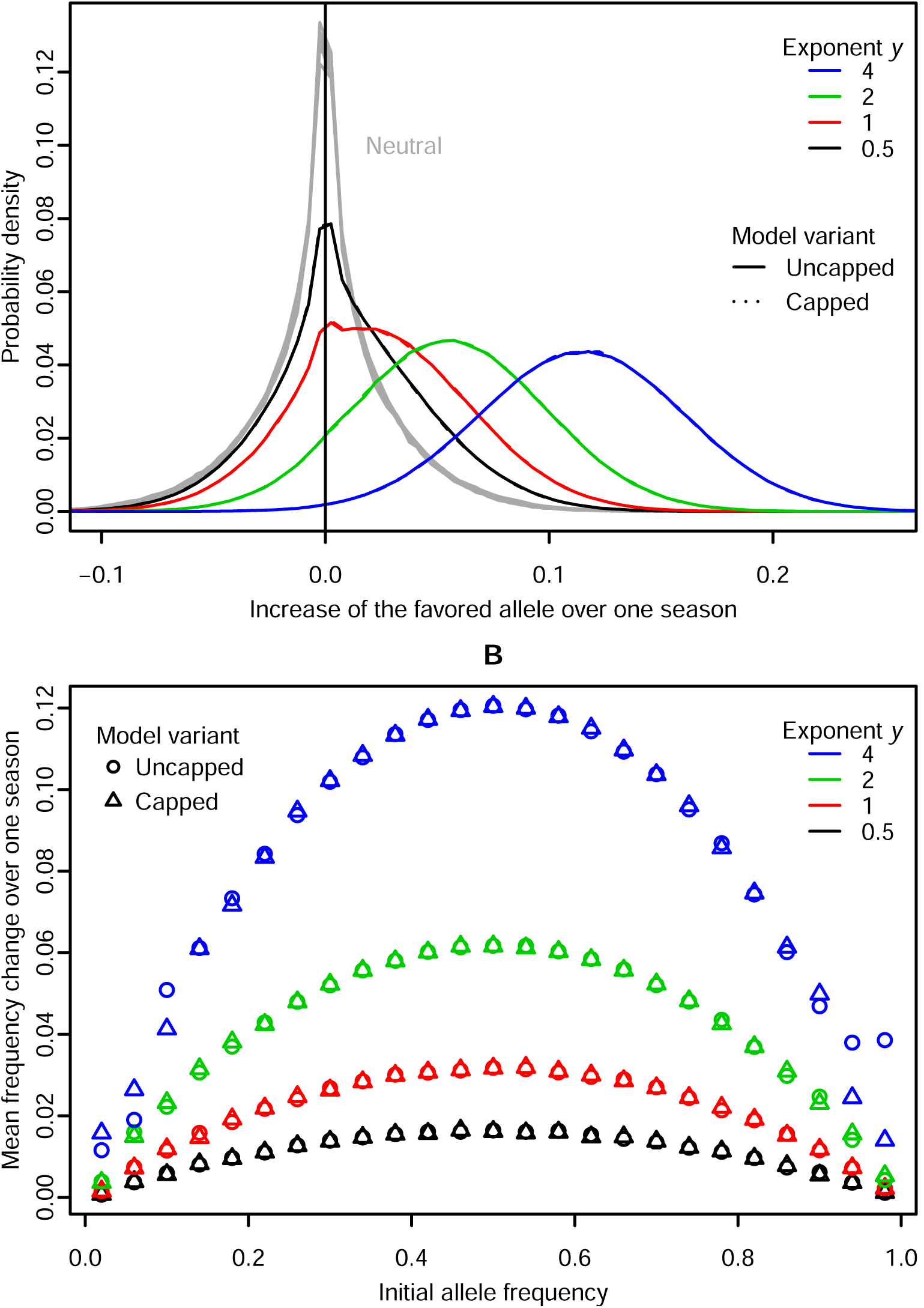
Detailed information on allele frequency change over one season for the scenarios with *d* = 0.7 in Fig. 9. (A) The distribution of the change in frequency for all loci over the currently favored allele over one season compared to neutrality (gray curve), and (B) the average frequency change of the currently favored allele over one season for *d* = 0.7 as a function of the frequency at the beginning of the season. *N* = 1000, *L* = 100, *g* = 15, *μ* = 10^−4^. In (A) the results for the capped vs. uncapped model variants are virtually indistinguishable.

**Figure S5:**
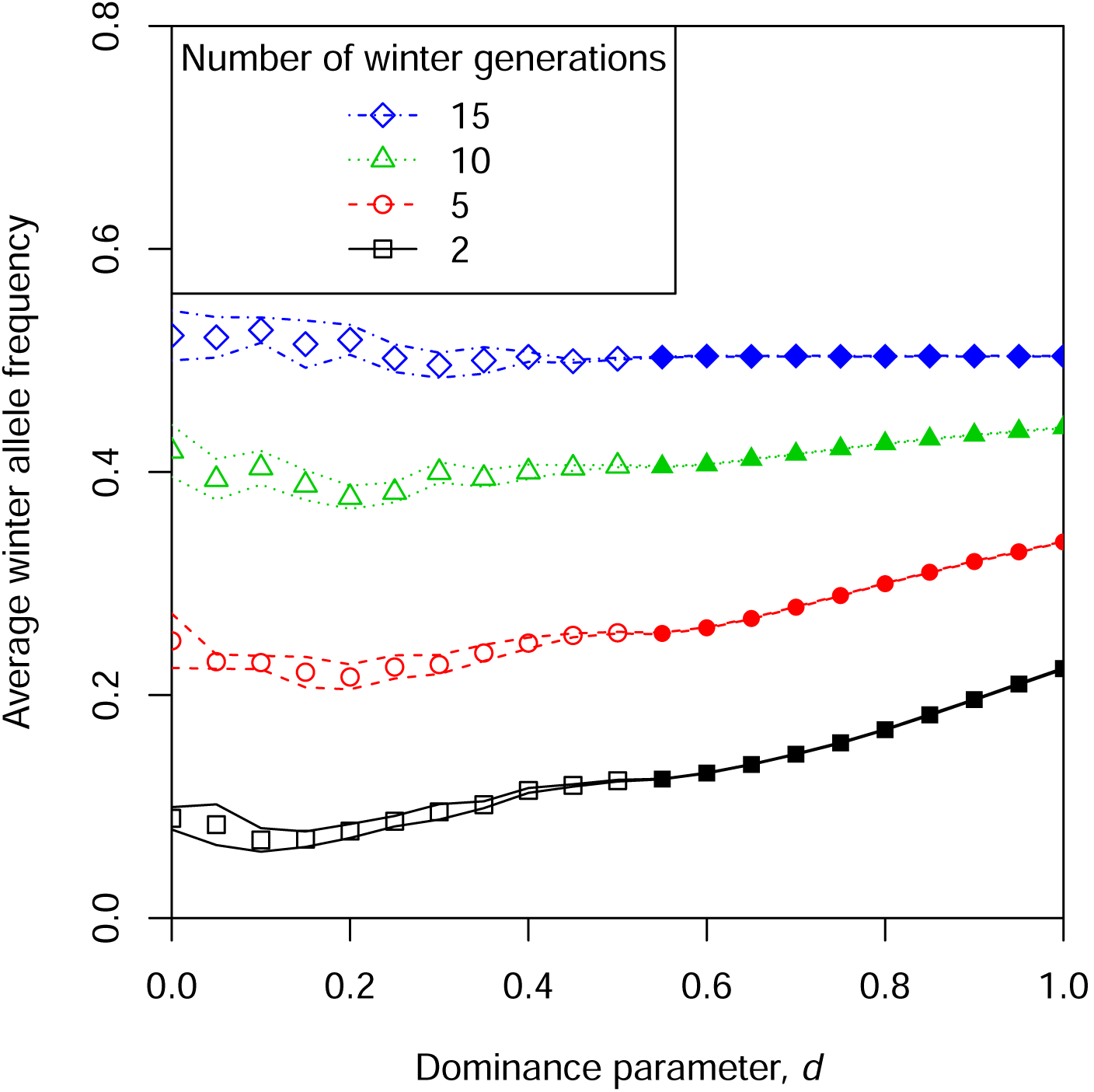
Influence of an imbalance in the number of generations in summer vs. winter. There are 15 generations of summer. The *y*-axis shows the winter allele frequency in the middle of winter, averaged across time and loci. Closed symbols represent parameter combinations that were classified as having stable polymorphism (see section Stability in asymmetric scenarios), open symbols represent unstable polymorphism. Lines indicate means ± two standard errors. Other parameters: *N* = 1000, *L* = 100, *y* = 4, *μ* = 10^−4^.

**Figure S6:**
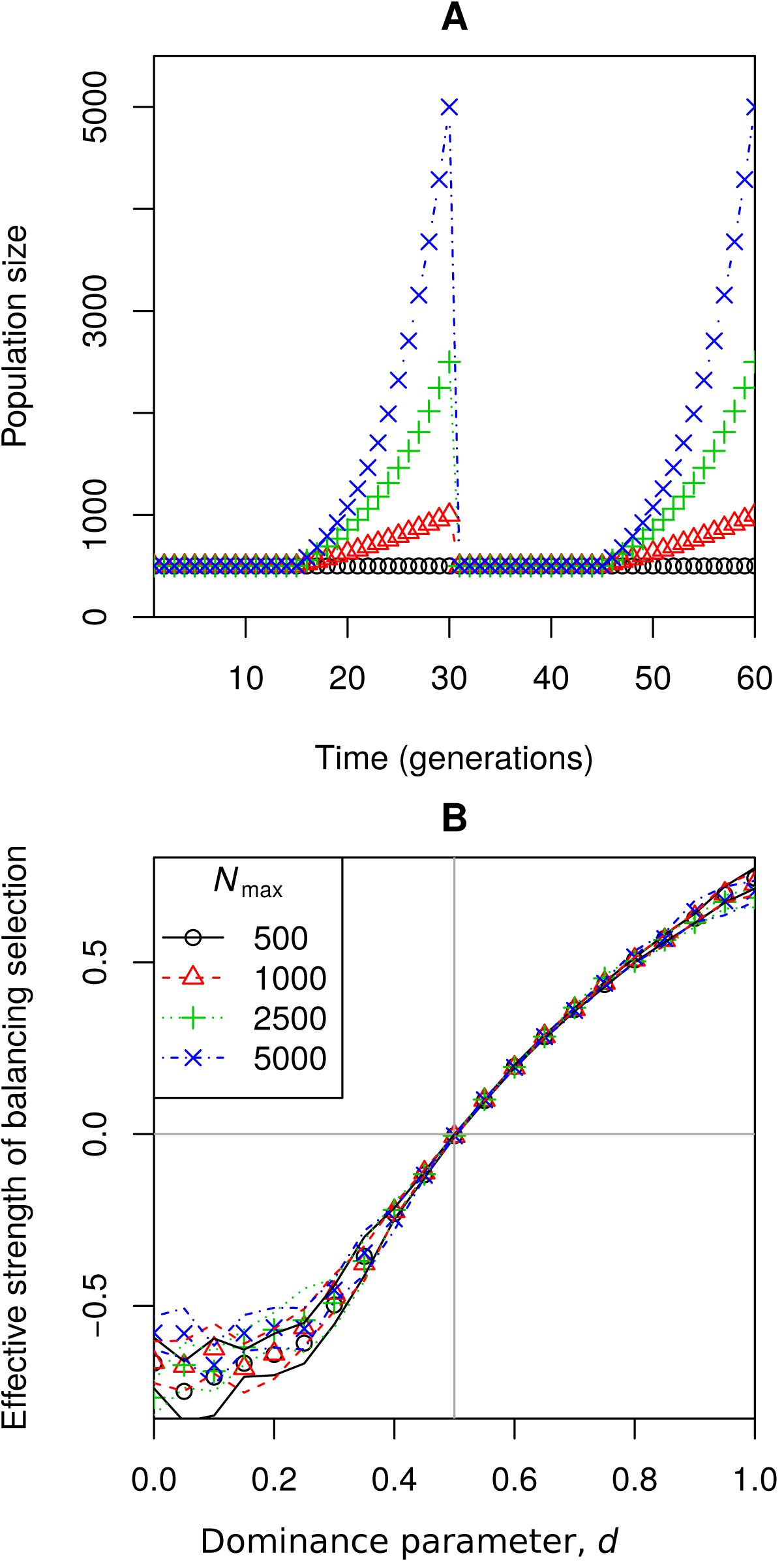
Robustness of results to seasonal changes in population size. (A) Four population-size trajectories. The trajectories are deterministic and cyclic and only two years are shown. Population size is 500 during winter and grows exponentially to different final sizes, *N*_max_, over summer (except for the case *N*_max_ = 500, where population size remains constant throughout the year). (B) Corresponding effective strength of balancing selection. *L* = 100, *g* = 15, *y* = 4, *μ* = 10^−4^.

**Figure S7:**
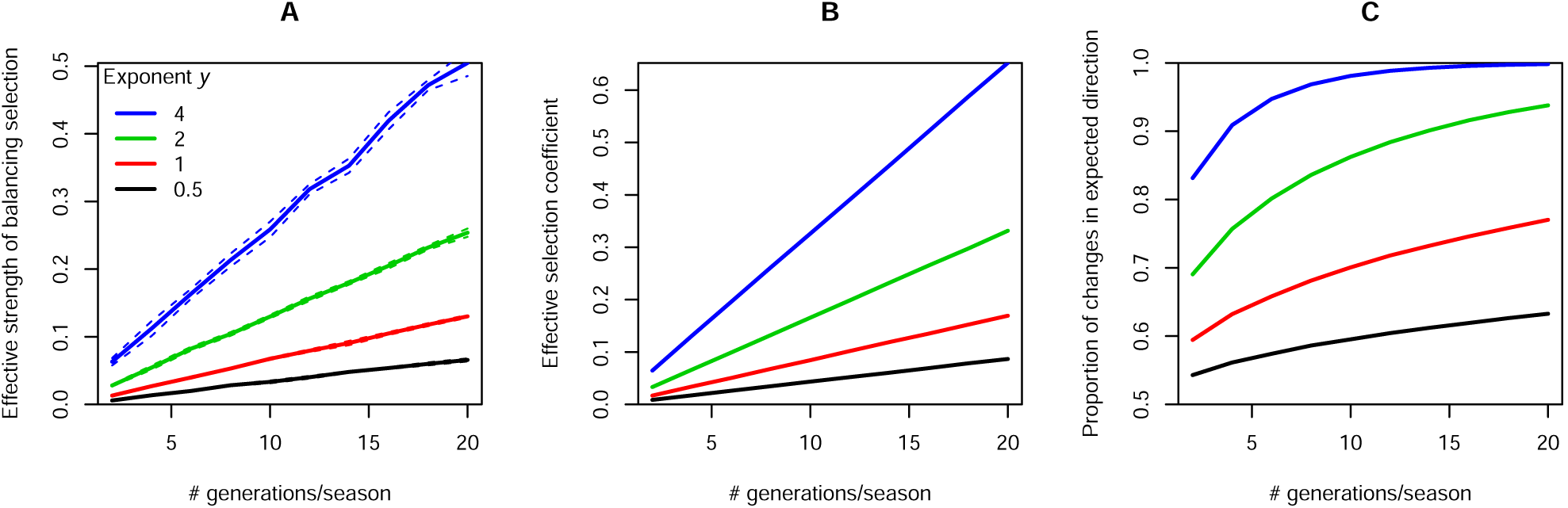
Influence of the number of generations per season, *g*, on (A) stability of polymorphism, (B) magnitude and (C) predictability of fluctuations. Dashed lines indicate means ± two standard errors, but they coincide with the solid lines in B and C. *N* = 1000, *L* = 100, *μ* = 10^−4^, *d* = 0.7.

**Figure S8:**
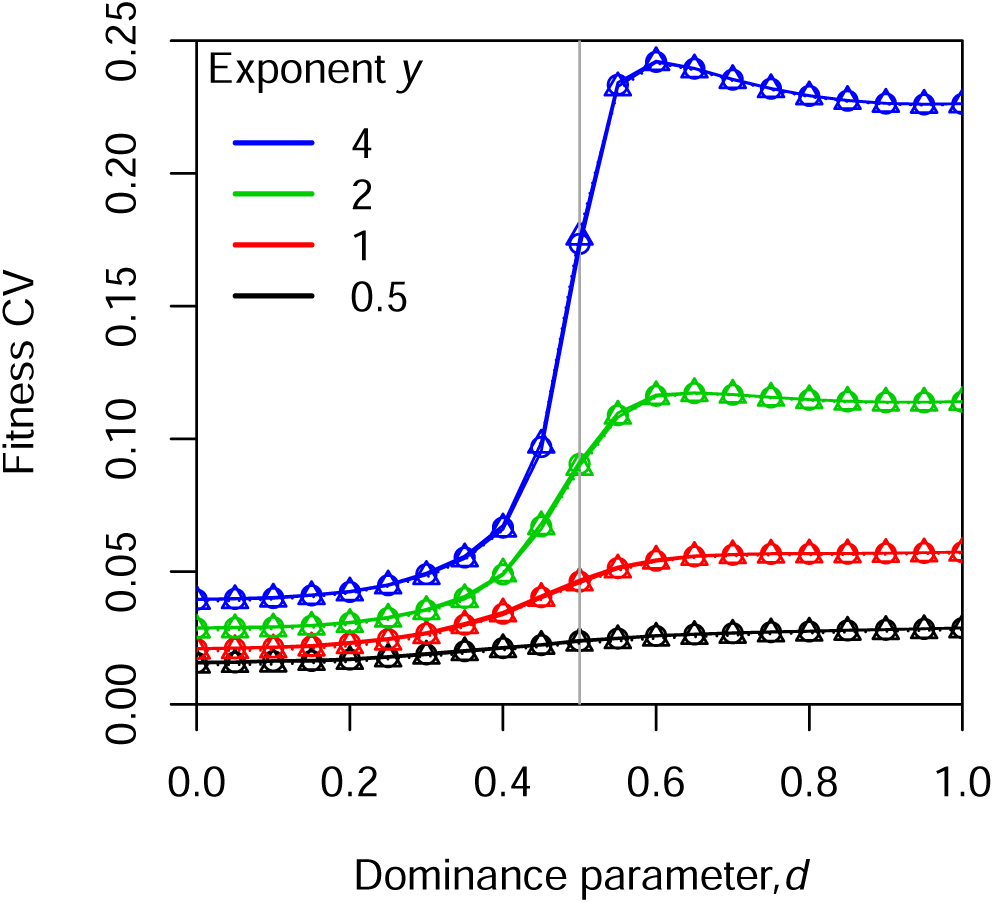
Coefficient of variation in fitness among individuals in the population as a function of the dominance parameter *d*. *N* = 1000, *L* = 100, *g* = 15, *μ* = 10^−4^. Symbols indicate the mean across replicates for the uncapped vs. capped model variant (often overlapping) and lines indicate means ± two standard errors for the uncapped case. Since the standard errors are so small, the lines are overlapping

**Figure S9:**
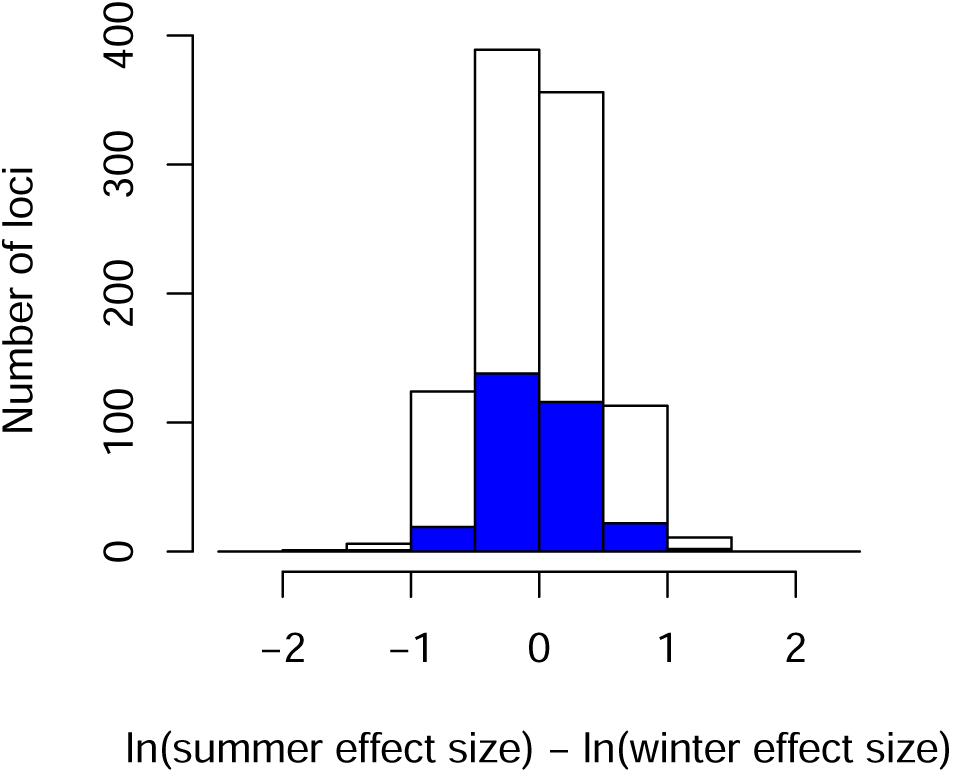
Histograms of the difference in the logarithms of seasonal effect sizes for stable (blue bars) compared to all polymorphisms (white bars) in Fig. 11 E, i.e. for *N* = 10, 000, *L* = 100, *g* = 10, *y* = 4, *μ* = 10^−4^.

